# GEMS – Enhancing Generalizable Binding Affinity Prediction by Removing Data Leakage and Integrating Language Model Embeddings into Graph Neural Networks

**DOI:** 10.1101/2024.12.09.627482

**Authors:** David Graber, Peter Stockinger, Fabian Meyer, Siddhartha Mishra, Claus Horn, Rebecca Buller

**Author notes:** shared senior authorship (Siddhartha Mishra, Claus Horn, Rebecca Buller).

## Abstract

The field of computational drug design requires accurate scoring functions to predict binding affinities for protein-ligand interactions. However, train-test data leakage between the PDBbind database and the CASF benchmark datasets has significantly inflated the performance metrics of currently available deep-learning-based binding affinity prediction models, leading to overestimation of their generalization capabilities. We address this issue by proposing PDBbind CleanSplit, a training dataset curated by a novel structure-based filtering algorithm that eliminates train-test data leakage as well as redundancies within the training set. Retraining current top-performing models on CleanSplit caused their benchmark performance to drop significantly, indicating that the performance of existing models is largely driven by data leakage. In contrast, our graph neural network model for efficient molecular scoring (GEMS) maintains high benchmark performance when trained on CleanSplit. Leveraging a sparse graph modeling of protein-ligand interactions and transfer learning from language models, GEMS is able to generalize to strictly independent test datasets.

---

Structure-based drug design (SBDD) aims to design small-molecule drugs that bind with high affinity to specific protein targets. In recent years, deep neural networks have begun to revolutionize the field, offering new possibilities for computational drug design. These include new protein folding models like RoseTTAFold All-Atom [1], AlphaFold3 [2] and Boltz-1 [3], which can also consider small-molecule ligands to predict potential binding conformations. Furthermore, generative artificial intelligence can design entirely novel protein-ligand interactions. For instance, RFdiffusion [4] can construct proteins around small-molecules starting from random clouds of amino acids while the denoising diffusion model DiffSBDD [5] generates novel ligands tailored to fit specific protein pockets. While these methods excel at generating diverse collections of protein-ligand interactions, these interactions are not necessarily characterized by drug-like affinity. Therefore, using these models for development of smallmolecule drugs requires *scoring functions* that can accurately predict the absolute binding affinities for protein-ligand poses and identify high-affinity complexes. Classical scoring functions, such as force-field-based, empirical, and knowledge-based methods implemented in docking tools like AutoDock Vina [6] and GOLD [7] are computationally intensive and show limited accuracy in binding affinity prediction [8–11]. Despite notable advancements in deep-learning-based scoring functions, including the design of many convolutional [12–23] and graph neural networks [24–32], accurately predicting binding affinities for protein-ligand poses remains an outstanding challenge.

In addition to the fact that many deep-learning-based scoring functions are either not publicly available or are difficult to implement, the key reason for their limited applicability currently is the observation that these scoring functions perform poorly, with considerably lower than expected accuracy on independent test data sets [33–36]. This large gap between benchmark and real-world performance has been attributed to the underlying training and evaluation procedures used for the design of these scoring functions. Typically, these models are trained on the PDBbind database [37], and their generalization is assessed using the Critical Assessment of Scoring Function (CASF) benchmark datasets [10]. However, several studies have reported a high degree of similarity between PDBbind and the CASF benchmarks. Due to this similarity, the performance on CASF overestimates the generalization capability of models trained on PDBbind [10, 38, 39]. Alarmingly, some of these models even perform comparably well on the CASF datasets after omitting all protein or ligand information from their input data. This suggests that the reported impressive performance of these models on the CASF benchmarks are not based on an understanding of protein-ligand interactions. Instead, memorization and exploitation of structural similarities between training and test complexes appear to be the main factors driving the observed benchmark performance of these models [35, 36, 40–42].

Our first goal in this paper is to further investigate the presence of a train-test data leakage between PDBbind and the commonly used CASF benchmarks. To this end, we propose a novel structure-based clustering algorithm to analyze and filter datasets of protein-ligand complex structures. By identifying large similarities between PBDbind and CASF datasets, our algorithm revealed a significant level of train-test data leakage. Going further, it also enabled us to devise a new split for the PBDbind dataset for providing a better setup for the training and testing of structure-based affinity prediction models. Our filtered training dataset, termed *PDBbind CleanSplit*, is strictly separated from the CASF benchmark datasets, turning them into true external datasets and enabling genuine evaluation of model generalizability.

To evaluate the true performance of recently published deep-learning-based scoring functions, we retrained the state-of-the-art binding affinity prediction models GenScore [30] and Pafnucy [12] on the PDBbind CleanSplit dataset with reduced data leakage. Although these models had previously shown excellent benchmark performance when trained on the original PDBbind dataset, their performance dropped markedly when trained on PDBbind CleanSplit, confirming that the prior high scores were largely driven by data leakage.

Recognizing that the generalization capability of existing deep-learning-based scoring functions might be much lower than previously thought, our second goal in this paper was to design a binding affinity prediction model with robustly validated generalization capability. To this end, we combined a novel graph neural network (GNN) architecture with transfer learning from large language models and trained this model on the filtered PDBbind CleanSplit. Despite the reduced data leakage, our Graph Neural Network for Efficient Molecular Scoring (GEMS) achieves state-of-the-art predictions on the CASF benchmark. Since all protein-ligand complexes that remotely resembled any from the CASF test set were excluded from training, we can confidently state that the performance of GEMS is not the result of exploiting data leakage, but genuinely reflects its capability to generalize to new complexes. Moreover, our ablation studies showed that GEMS fails to produce accurate predictions when protein nodes are omitted from the graph, suggesting that its predictions are based on a genuine understanding of protein-ligand interactions.

GEMS is a promising tool with broad potential impact on the field of structure-based drug design (SBDD). Generative models like RFdiffusion and DiffSBDD can generate libraries of novel protein-ligand interactions, but their potential in drug design has been bottlenecked by the lack of accurate models to predict binding affinities for these interactions. GEMS fills this critical gap in SBDD. With its robust generalization capabilities evaluated on strictly independent datasets, it provides the prediction accuracy needed to identify interactions with therapeutic potential. To enable researchers to leverage and further develop GEMS, we have made all Python code publicly available in an easy-to-use format.

## Results

### PDBbind Dataset Filtering

To gain the ability to identify and remove structural similarities in datasets of protein-ligand complexes, we set out to design a structure-based clustering algorithm (**Fig. 1**). In this algorithm, the computation of similarity between two protein-ligand complexes is based on a combined assessment of protein similarity (TM-scores [43]), ligand similarity (Tanimoto scores [44]), and binding conformation similarity (pocket-aligned ligand RMSD). Combining these three metrics allows a robust and detailed comparison of protein-ligand complex structures. Importantly, in difference to traditional sequence-based analysis approaches, our multi-modal filtering can identify complexes with similar interaction patterns, even when the proteins have low sequence identity (Supplementary Fig. 1).

**Figure 1:**
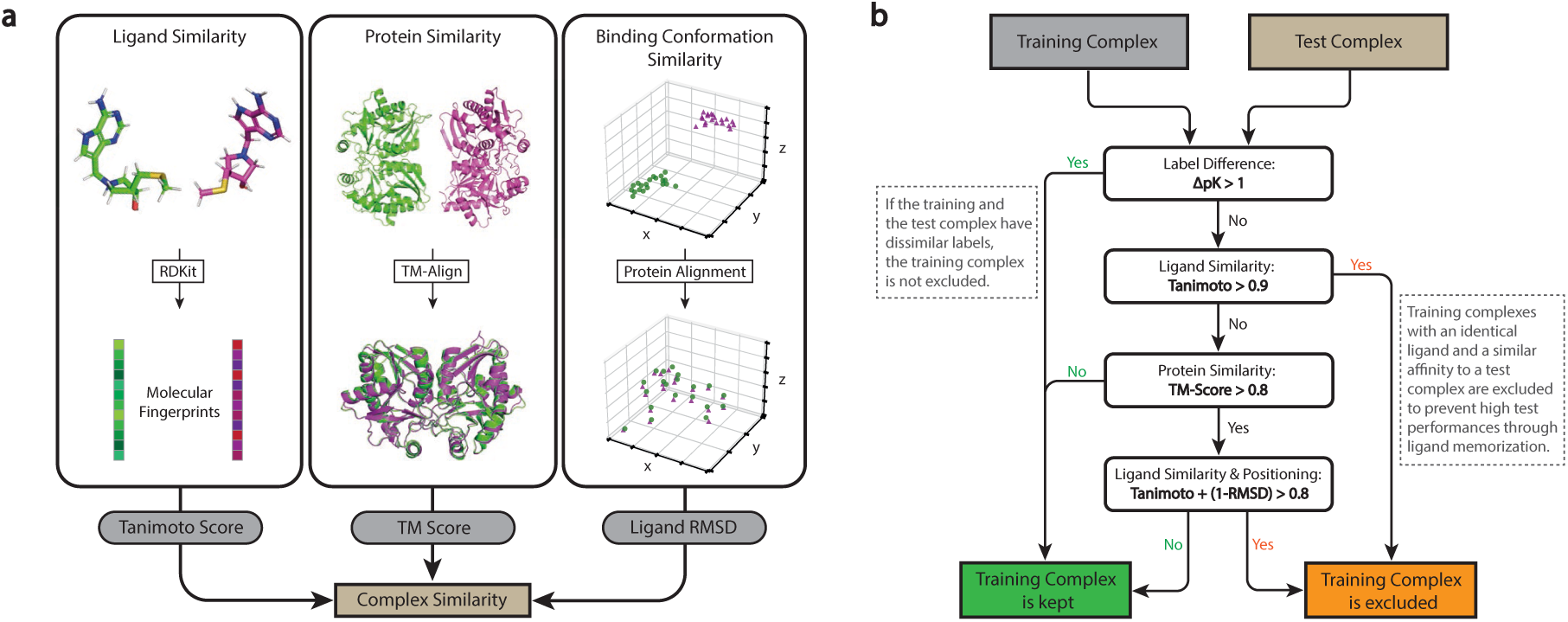
Overview of the Similarity Computation Between Two Protein-Ligand Complexes: **a)** Our structure-based dataset filtering algorithm evaluates structural similarity of protein-ligand complexes using a combination of protein similarity (TM-scores), ligand similarity (Tanimoto scores) and binding conformation similarity (pocket-aligned ligand RMSD). The Tanimoto scores identify chemically similar ligands and range from 0 (no similarity) to 1 (identical). TM-scores are computed with TM-align, a tool that compares protein structures by finding the optimal alignment of their three-dimensional structures and outputs a score ranging from 0 (no similarity) to 1 (identical). This score identifies proteins with high structural similarity, even when sequence identity is low (e.g. when one protein is a substructure of the other). Pocket-aligned ligand RMSD scores compare the positioning of ligands within aligned protein pockets. Ligands are transformed into the same coordinate frame using the optimal alignment from TM-align, and a root-mean-square-deviation (RMSD) calculation provides a quantitative measure of positional similarity. **b)** Decision tree showing the exclusion criteria and the information flow of the filtering algorithm when comparing a training and a test complex. The first layer of the algorithm compares the affinity labels of the complexes (pK values, see Methods). Training complexes with high structural similarity but different activity are not excluded, to avoid excluding data points that can provide valuable insights into activity cliffs. The second layer excludes all training complexes with similar affinity and identical ligand (Tanimoto *>* 0.9) to make the test complex ligands unique and avoid successful predictions through ligand memorization. The third and fourth layer exclude training complexes based on protein similarity and a combined assessment of ligand and binding conformation similarity. This four-layer approach identifies complexes with similar interaction patterns, even when traditional sequence-based methods would overlook these similarities.

By comparing all CASF complexes to all PDBbind complexes, we identified a large number of train-test pairs with exceptionally high similarity, sharing not only similar ligand and protein structures but also comparable ligand positioning within the protein pocket, unsurprisingly also being accompanied by closely matched affinity labels (**Fig. 2a**). Consequently, these structures provide nearly identical input data points to the model, enabling accurate prediction of test data point labels through simple memorization. According to the thresholds of our filtering algorithm, almost 600 PDBbind training complexes were detected to share such similarities with a CASF complex, involving 49% of all CASF complexes. These findings reveal a clear train-test data leakage when models are trained on PDBbind and tested on the CASF benchmark datasets, with nearly half of the CASF complexes not presenting novel challenges to these models.

**Figure 2:**
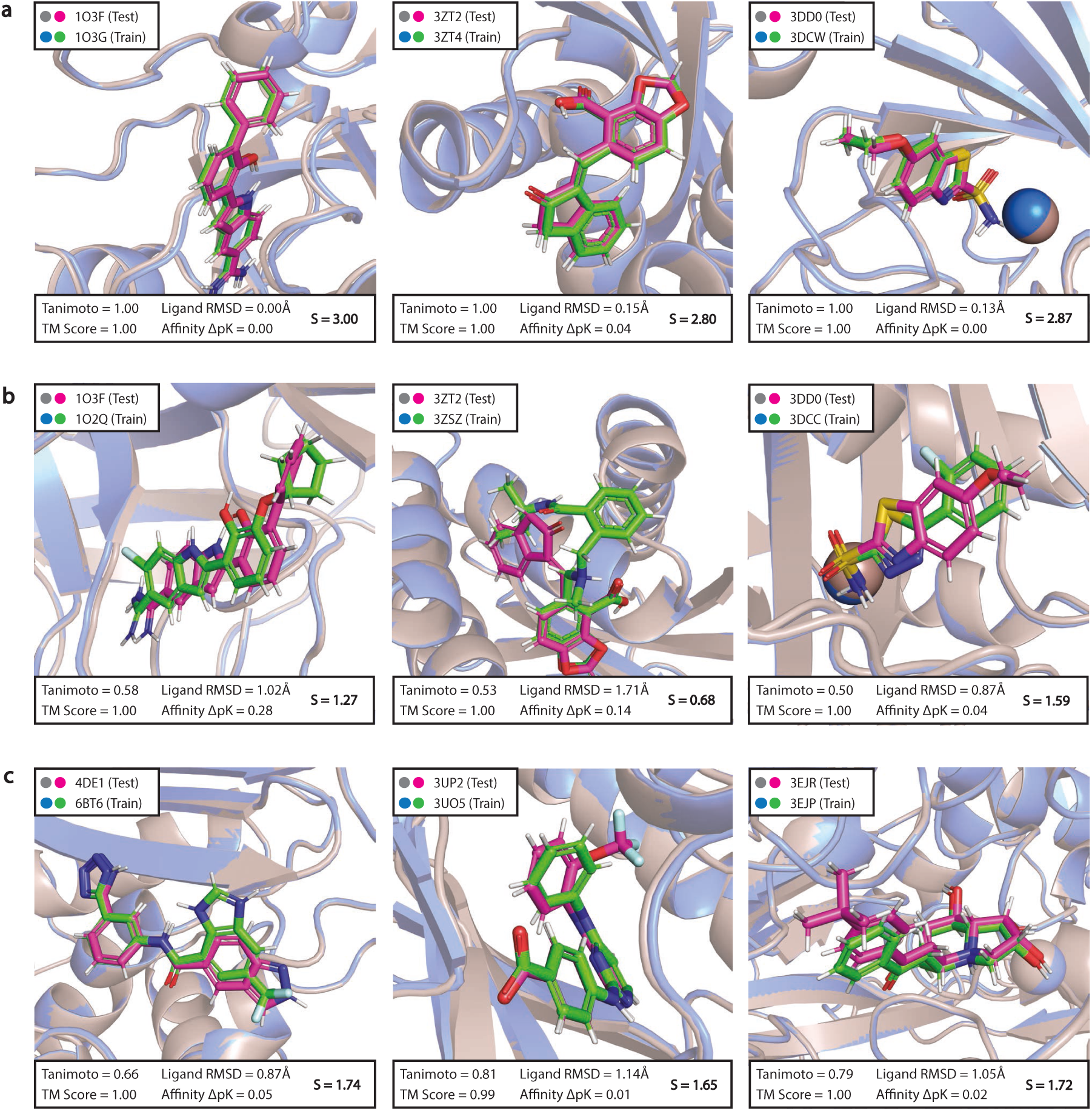
Superpositions of Complexes Highlighting Train-Test Structural Similarities Before and After Filtering: **a)** Superpositions of the most prominent train-test similarities before applying the filtering algorithm. **b)** Superpositions of the same test complexes as in a), now shown with the most similar training complexes found in PDB CleanSplit. **c)** Superpositions of the closest train-test similarities that remained post-filtering in the dataset PDBbind CleanSplit. Protein structures from the test and training datasets are depicted as grey and blue cartoons, respectively, with ligands shown in magenta (test) and green (train). Below each superposition, the Tanimoto score, TM-score, ligand RMSD, and affinity difference (ΔpK) is shown, which are combined into an overall similarity score *S* = TM-Score + Tanimoto + (1 *−* Ligand RMSD) *−* ΔpK. This S-score was computed for all possible train-test pairs and served to select representative complexes for this figure. For each training dataset, the depicted pairs were selected from the top 5 pairs with the highest *S*, prioritizing good ligand visibility. In some superpositions (1O3F/1O3G and 3DD0/3DWC), one structure has been slightly shifted to enhance visibility of structural differences. Original PyMol sessions of all of superpositions are provided on GitHub.

Our filtering algorithm reduced train-test data leakage by excluding all training complexes that closely resemble any CASF test complex (**Fig. 1b**). Additionally, it removed all training complexes with ligands identical to those in the CASF test complex (Tanimoto *>* 0.9), ensuring that the ligands in the test datasets are never encountered with similar affinity during model training. This step provides an additional safeguard against ligand-based data leakage, addressing prior research that shows GNNs for binding affinity predictions often rely on ligand memorization to make affinity predictions [40]. Together, this filtering excluded 4.5% of all training complexes. The remaining train-test pairs with the highest similarity after filtering exhibited clear structural differences (**Fig. 2c**), highlighting the effectiveness of our filtering algorithm in removing structurally similar data points. The resulting filtered training dataset is strictly separated from the CASF datasets, allowing models trained on it to be evaluated on the CASF benchmark, offering a genuine assessment of their generalization to unseen protein-ligand complexes.

In addition to the train-test overlap, we found significant similarity clusters within the training dataset itself. According to the thresholds of our filtering algorithm, nearly 50% of all training complexes are part of a similarity cluster. This means that random splitting inadvertently leads to inflated validation performance metrics, as some validation complexes can be predicted by matching labels with similar training complexes. Consequently, it is not surprising that models trained on a dataset with such extensive redundancies perform structure-matching, thereby settling for an easily attainable local minimum in the loss landscape. We hypothesized that this redundancy hampers model generalization, as it encourages memorization, leading to models that rely on exploiting structural similarities. Thus, we proposed that binding affinity prediction models would benefit from a more diverse dataset as a robust basis for training. To test this hypothesis, our filtering algorithm includes a step to reduce training dataset redundancy. To find an optimal trade-off between maximizing dataset size and minimizing redundancy, we used adapted filtering thresholds to identify and eliminate the most striking similarity clusters (see Methods). Using these adapted thresholds, our filtering algorithm iteratively removed complexes from the training dataset until all similarity clusters were resolved. Ultimately, this process resulted in the removal of 7.5% of all training complexes.

Given the extensive data leakage between PDBbind and the CASF benchmarks, the performance reported by many published models trained on these datasets likely overestimate their true generalization capabilities. Combining our strategies for reducing train-test data leakage and minimizing training dataset redundancy (see Method section: Filtering Algorithm), we have created a new refined split of the PDBbind dataset, which we call PDBbind CleanSplit.

### Search Algorithms

To illustrate the effect of train-test data leakage on model performance, we devised a simple algorithm that predicts the affinity of each CASF test complex by identifying the five most similar training complexes and averaging their affinity labels. This algorithm showed competitive CASF2016 prediction root-mean-square-error (RMSE) compared to some published deep-learningbased scoring functions (Pearson R=0.716, RMSE=1.517). To explore whether ligand memorization alone is sufficient for accurate CASF predictions, we modified the algorithm to search for the five training complexes with the most similar ligands. Averaging the labels of these complexes produced similarly high performance (Pearson R=0.707, RMSE=1.539), confirming prior research on the importance of ligand memorization [40] (**Table 1**).

**Table 1:**
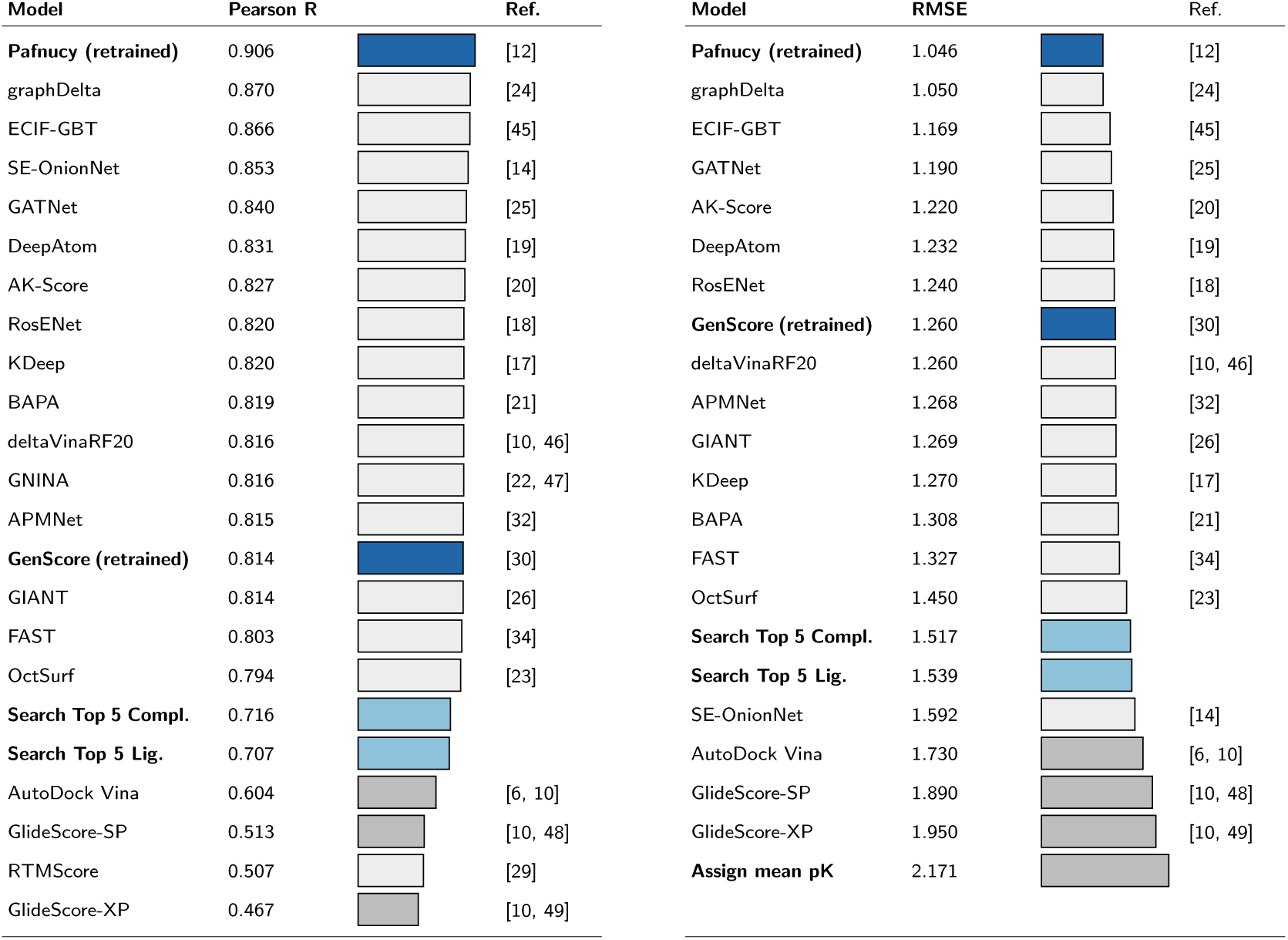
Comparison of Model Performance: Reported CASF2016 scoring performance (Pearson correlation coefficients and RMSE values) for all published deep-learning-based scoring functions that to our knowledge have evaluated their scoring performance on the complete CASF2016 (n=285) dataset. Models with blue bars and bold text have been generated or trained in this study (Pafnucy, GenScore and our two search algorithms “Search Top 5 Complexes” and “Search Top 5 Ligands”). Models with light-grey bars are published deep-learning-based binding affinity prediction models, with performance values taken from literature. Models with dark-grey bars are classical scoring functions (AutoDock Vina, GlideScore). As a baseline, the lowest bar in the RMSE table shows the RMSE that is achieved when the average training dataset label is assigned to all CASF2016 complexes.

| Model | Pearson R |  | Ref. | Model | RMSE |  | Ref. |
| --- | --- | --- | --- | --- | --- | --- | --- |
| **Pafnucy (retrained)** | 0.906 |  | [12] | **Pafnucy (retrained)** | 1.046 |  | [12] |
| graphDelta | 0.870 |  | [24] | graphDelta | 1.050 |  | [24] |
| ECIF-GBT | 0.866 |  | [45] | ECIF-GBT | 1.169 |  | [45] |
| SE-OnionNet | 0.853 |  | [14] | GATNet | 1.190 |  | [25] |
| GATNet | 0.840 |  | [25] | AK-Score | 1.220 |  | [20] |
| DeepAtom | 0.831 |  | [19] | DeepAtom | 1.232 |  | [19] |
| AK-Score | 0.827 |  | [20] | RosENet | 1.240 |  | [18] |
| RosENet | 0.820 |  | [18] | **GenScore (retrained)** | 1.260 |  | [30] |
| KDeep | 0.820 |  | [17] | deltaVinaRF20 | 1.260 |  | [10, 46] |
| BAPA | 0.819 |  | [21] | APMNet | 1.268 |  | [32] |
| deltaVinaRF20 | 0.816 |  | [10, 46] | GIANT | 1.269 |  | [26] |
| GNINA | 0.816 |  | [22, 47] | KDeep | 1.270 |  | [17] |
| APMNet | 0.815 |  | [32] | BAPA | 1.308 |  | [21] |
| **GenScore (retrained)** | 0.814 |  | [30] | FAST | 1.327 |  | [34] |
| GIANT | 0.814 |  | [26] | OctSurf | 1.450 |  | [23] |
| FAST | 0.803 |  | [34] | **Search Top 5 Compl.** | 1.517 |  |  |
| OctSurf | 0.794 |  | [23] | **Search Top 5 Lig.** | 1.539 |  |  |
| **Search Top 5 Compl.** | 0.716 |  |  | SE-OnionNet | 1.592 |  | [14] |
| **Search Top 5 Lig.** | 0.707 |  |  | AutoDock Vina | 1.730 |  | [6, 10] |
| AutoDock Vina | 0.604 |  | [6, 10] | GlideScore-SP | 1.890 |  | [10, 48] |
| GlideScore-SP | 0.513 |  | [10, 48] | GlideScore-XP | 1.950 |  | [10, 49] |
| RTMScore | 0.507 |  | [29] | **Assign mean pK** | 2.171 |  |  |
| GlideScore-XP | 0.467 |  | [10, 49] |  |  |  |  |

When using the two search algorithms on the filtered PDBbind CleanSplit, we found that this led to a dramatic drop in their CASF prediction performance. Averaging the affinity label of the five most similar training complexes resulted in an RMSE of 1.648 and a Pearson correlation of 0.653. Using the labels of the five complexes with the most similar ligands resulted in an RMSE of 1.711 and a Pearson correlation of 0.625 (**Fig. 3a,b**).

**Figure 3:**
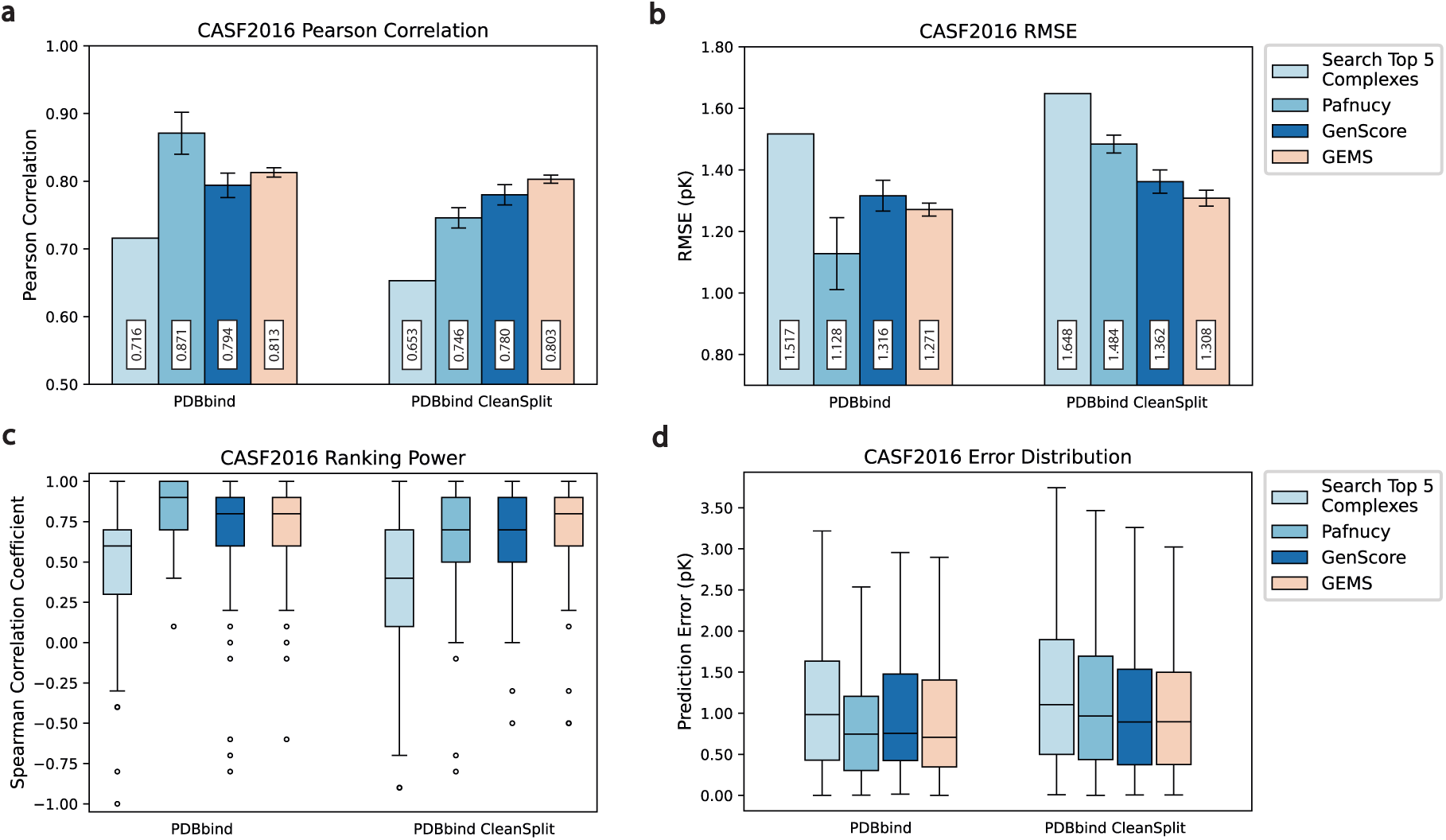
Prediction Accuracy of GEMS, Pafnucy and GenScore in Dependence of Dataset Filtering: **a)** CASF2016 Pearson correlation coefficients of a simple search algorithm (Search Top 5 Complexes), Pafnucy, GenScore, and GEMS when trained on the original PDBbind dataset (left) and the filtered PDBbind CleanSplit dataset (right). **b)** CASF2016 prediction RMSE values of a simple search algorithm (Search Top 5 Complexes), Pafnucy, GenScore and GEMS when trained on the original PDBbind dataset (left) and the filtered PDBbind CleanSplit dataset (right). **c)** Ranking power: The distribution of Spearman correlation coefficients for the search algorithm, Pafnucy, GenScore and GEMS across all 57 clusters of CASF2016, presented separately for training on PDBbind (left) and PDBbind CleanSplit (right). **d)** Distribution of prediction errors (pK unit) across all CASF2016 complexes shown for a simple search algorithm (Search Top 5 Complexes), Pafnucy, GenScore, and GEMS when trained on the original PDBbind dataset (left) and the filtered PDBbind CleanSplit dataset (right). For each Pafnucy, GenScore and GEMS, five models were trained with five different train-validation splits. The performance values in **a)** and **b)** reflect the mean CASF2016 performance of the five models, with error bars representing the standard deviation across the performance of the five models. To compute the distributions of the Spearman correlation coefficients and prediction errors in **c)** and **d)**, the predictions of the five models were averaged into an ensemble prediction.

Overall, the success of these two search algorithms on the unfiltered PDBbind illustrates how much training data memorization can boost CASF prediction accuracy when training on PDBbind. The low accuracy of the same algorithms on PDBbind CleanSplit, however, demonstrates that the train-test similarities have been largely removed through our filtering. For models trained on PDBbind CleanSplit, training data memorization is not sufficient for high CASF performance.

### Retraining State-Of-The-Art Models

Building on the findings we obtained with our simple search algorithms, we set out to investigate whether the train-test data leakage within PDBbind has similarly inflated the benchmark performance of published state-of-the-art models. Toward this goal, we began retraining the top-performing models to reproduce their reported results and evaluate their performance when trained on PDBbind CleanSplit. Unfortunately, we encountered significant obstacles that made reproducing most of these models virtually impossible. These obstacles included the absence of public code repositories, the availability of code that supported inference only, a lack of training instructions, and reliance on refined, augmented, or proprietary datasets that are not publicly accessible. However, retraining was successful for two models, GenScore [30] and the well-known Pafnucy binding affinity prediction model [12]. Pafnucy, published in 2018, was originally trained on the 2016 version of PDBbind, and its benchmark performance has been surpassed by newer models over time. We retrained Pafnucy using a 5-fold cross-validation approach on the more recent 2020 version of PDBbind according to the authors’ instructions. This increased the CASF performance of Pafnucy to an RMSE of 1.046, making it the best-performing binding affinity prediction model among all models that have, to our knowledge, been evaluated on the complete CASF2016 dataset to date (**Table 1**). As a second test case, we retrained GenScore [30], a model which also reports state-of-the-art performance in absolute binding affinity prediction.

As a next step in our evaluation, we repeated the Pafnucy training using our PDBbind CleanSplit dataset. Strikingly, the performance of the newly trained Pafnucy model on the CASF2016 benchmark dropped substantially, approaching the level of the simple search algorithms (**Fig. 3**). GenScore proved to be more robust, as it suffered a much smaller relative drop in scoring performance upon switching the training dataset to PDBbind CleanSplit (**Fig. 3**) These performance differences support our hypothesis that the reported performance of many published binding affinity prediction models are boosted by data leakage and that the true generalization capabilities of many models likely are much lower than reported.

### GNN for Efficient Molecular Scoring (GEMS)

Aiming to create a scoring function that better generalizes to new data, we developed GEMS, a graph-based model for proteinligand binding affinity prediction. GEMS models protein-ligand structures as interaction graphs enhanced with embeddings from language models and processes these graphs through a series of graph convolutions to predict binding affinities (**Fig. 4**).

**Figure 4:**
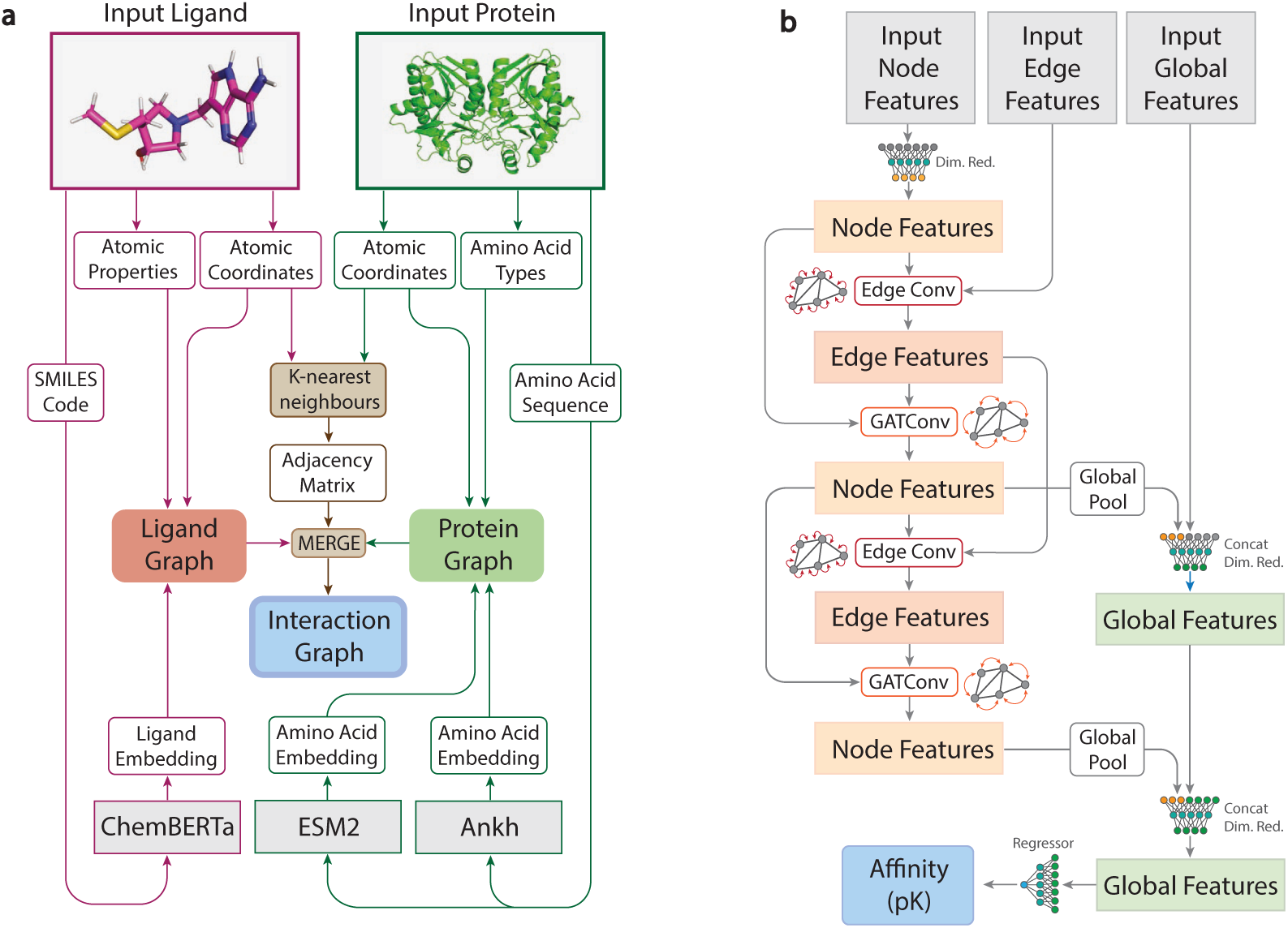
Graph-Based Modeling of Protein-Ligand Interactions and GEMS Model Architecture.: **a)** Schematic overview of the graph construction process used to model protein-ligand complexes in a sparse, rotation- and translation-invariant graph representation enhanced with language model embeddings. The core of these graph representations consists of an atom-level molecular graph of the ligand molecule (magenta) combined with an amino acid-level graph of the protein pocket (green). During the merging of ligand and protein graphs, additional edges are introduced to connect ligand graph nodes (atoms) to protein graph nodes (amino acids) based on spatial proximity between ligand atoms and the atoms of the amino acids (interaction distance of 5Å), computed using a K-nearest neighbors algorithm. The amino acid nodes are featurized with their type and embeddings derived from the protein language models ESM2 [50] and Ankh [51]. The ligand graph is featurized with atomic properties. The global features are intialized with a ligand embedding from the language model ChemBERTa-2 [52]. **b)** GEMS model architecture for processing interaction graphs composed of node features, edge features, and global context features. After an initial node feature dimensionality reduction (Dim.Red.), node and edge features are transformed through an alternating sequence of node convolutions (GATConv) and edge convolutions (EdgeConv). The global graph features are dynamically updated throughout this process, incorporating pooled node representations after each node convolution. A final pK value prediction is made from the updated global features using a fully connected neural network.

When trained on PDBbind, all tested model architectures showed benchmark performance comparable to those of the top deep-learning-based scoring functions reported to date. Training these models on PDBbind CleanSplit initially resulted in much lower benchmark performance, as would be expected when removing data leakage. However, after substantial architectural optimizations and the integration of language model embeddings to enrich the feature space of the graphs, our GEMS model achieved competitive results on the CASF2016 benchmark (**Fig. 3**). With a prediction RMSE of 1.308 and a Pearson Correlation of 0.803, GEMS considerably outperforms Pafnucy (RMSE=1.484, Pearson R=0.746) and GenScore (RMSE=1.362, Pearson R=0.780) when trained on PDBbind CleanSplit (**Fig. 3**), demonstrating its ability to generalize to unseen complexes. Moreover, GEMS even surpasses the reported performance metrics of several other deep-learning-based scoring functions that benefited from extensive train-test data leakage affecting 49% of all test data points (**Table 1**).

In addition to its scoring power, we also evaluated GEMS’ ranking power and its performance in different protein families. The CASF2016 dataset features 57 clusters, each consisting of five identical proteins paired with diverse ligands that span a wide range of binding affinities **(Supplementary Table 1)**. The ranking power of a scoring function refers to its ability to accurately rank ligands for a given target protein based on their binding affinities. When trained on PDBbind CleanSplit, GEMS outperformed both Pafnucy and GenScore, as underscored by the distribution of Spearman correlation coefficients across all 57 CASF2016 clusters (**Fig. 3c**). Moreover, GEMS showed high Pearson correlation coefficients and favourable absolute error distributions across most clusters (Extended Data Fig. 1).

#### Ablation

When trained on the original PDBbind, all tested GEMS model variants achieved competitive CASF2016 performance even after removing all protein information from the input data (RMSE=1.424) (**Fig. 5a**, Extended Data Fig. 4a). According to this evaluation, these models - when trained only on ligands - make more accurate predictions than the scoring function of AutoDock Vina and even outperform several published deep-learning-based scoring functions. These results align with other studies indicating that models trained on PDBbind can achieve remarkably high performance on the CASF benchmark even when one interaction partner is deleted from the input data [35, 36, 41, 42]. As these models received no protein information, their predictions are clearly not based on an understanding of protein-ligand interactions.

**Figure 5:**
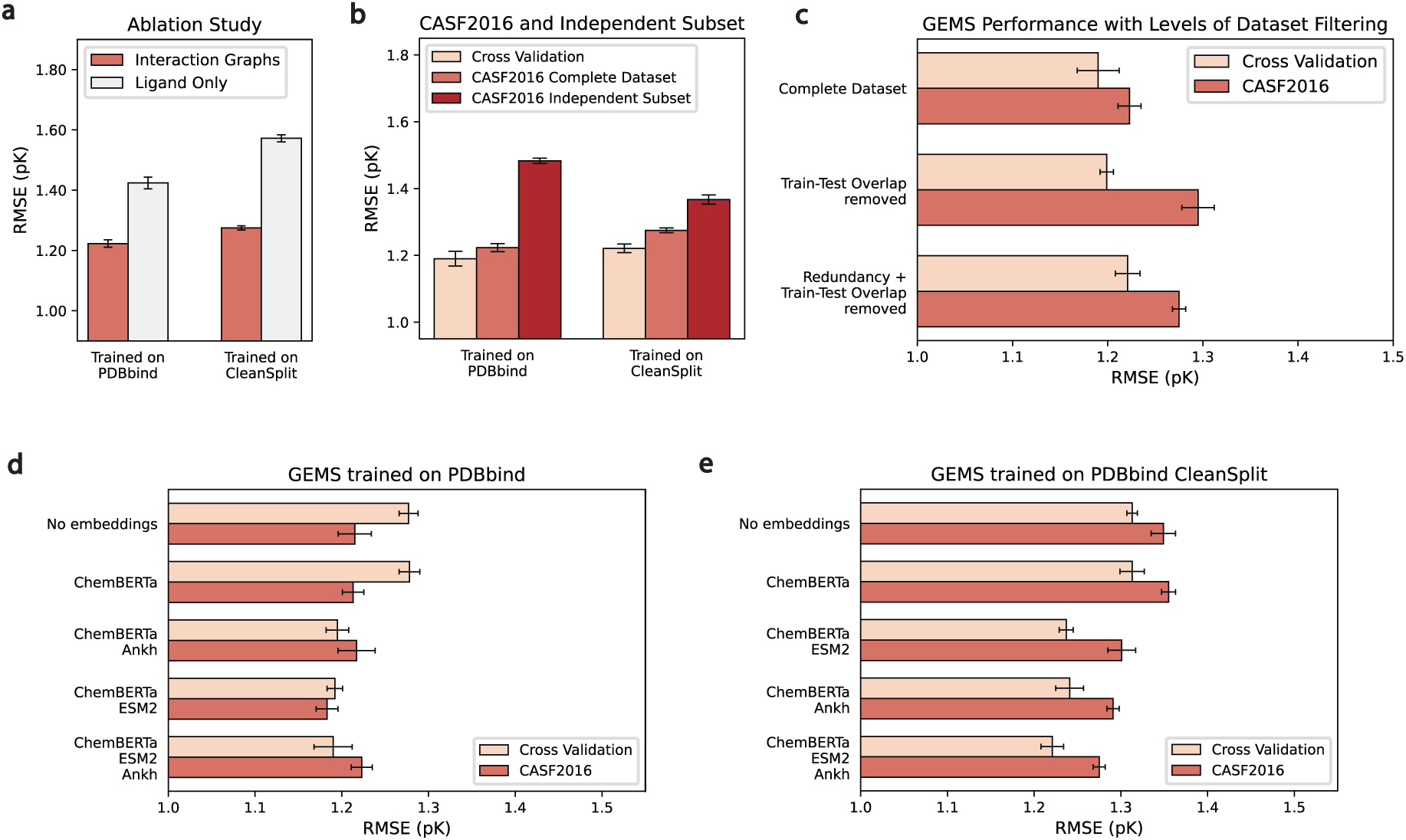
Impact of Ablation and Dataset Filtering on GEMS Performance: **a)** Ablation study showing the impact of removing all protein information from the graphs on CASF2016 (n=285) performance, comparing GEMS models trained on PDBbind (left) and PDBbind CleanSplit (right). **b)** Comparison of GEMS performance on cross-validation (CV), CASF2016, and the independent subset of CASF2016 (n=155) for models trained on PDBbind (left) and PDBbind CleanSplit (right). **c)** CASF2016 performance of GEMS with varying levels of training dataset filtering: complete dataset (PDBbind), train-test overlap removed, and both overlap and redundancy removed (PDBbind CleanSplit). **d)** Effect of incorporating language model embeddings on the cross-validation and CASF2016 (n=285) performance of GEMS models trained on the original PDBbind dataset. **e)** Effect of incorporating language model embeddings on the cross-validation and CASF2016 (n=285) performance of GEMS models trained on the PDBbind CleanSplit. Performance values represent the mean values of five models trained with 5-fold cross-validation at different random seeds. Error bars represent data uncertainty, calculated as the standard deviation of the performance across the five models.

In contrast, when GEMS was trained on PDBbind CleanSplit, it produced very inaccurate benchmark predictions once protein nodes were omitted from the input data (RMSE = 1.572). This significant performance drop indicates that when data leakage and redundancies are reduced, models must rely on an understanding of protein-ligand interactions to make accurate predictions.

#### Generalization to independent subset of CASF dataset

To explore whether models trained on PDBbind CleanSplit can generalize better to unseen complexes, we evaluated the performance of our models on a subset of the CASF2016 benchmark dataset, which is independent even before filtering the training data. According to the stringent similarity thresholds of our filtering algorithm, a fraction of the CASF2016 test dataset (143/285 complexes) is independent, with no similar complexes present in PDBbind. This independent subset thus provides a more reliable measure of generalization capability for models trained on PDB-bind. While GEMS trained on PDBbind showed high overall performance on the complete CASF2016 dataset (RMSE=1.223), the performance on the independent subset is notably lower (RMSE=1.483). Notably, when tested on the same independent subset, the GEMS model trained on PDBbind CleanSplit performed better than the model trained on PDBbind (RMSE=1.367) despite the significant training dataset size reduction, suggesting that the true generalization capability of the models trained on PDBbind CleanSplit is indeed superior (**Fig. 5b**, Extended Data Fig. 4b).

#### Influence of training redundancy

Our filtering approach for compiling PDBbind CleanSplit not only reduced train-test overlap but also training dataset redundancies. This reduction led to a decrease in validation performance, supporting our hypothesis that these redundancies had previously inflated validation results. While the reduction of the train-test overlap led to the anticipated drop in test set performance, eliminating the redundancies from the training dataset improved performance again (**Fig. 5c**, Extended Data Fig. 4c). While this process removed 1451 training complexes, which would conventionally be anticipated to reduce model performance, we observed a beneficial impact of redundancy removal across all replicates of the tested model variants. This positive effect suggests that extensive redundancies can be problematic in affinity prediction models, as models are prone to overfit to these clusters to minimize the training loss. Such overfitting interferes with learning the causal relationships behind molecular binding affinity and leads to models with lower generalization. In this case, distilling the training data to a smaller but more diverse collection of the most relevant complexes can be helpful for model training.

#### Influence of language model embeddings

Models trained on PDBbind and PDBbind CleanSplit exhibit a distinct response to the incorporation of language model embeddings (**Fig. 5d,e**, Extended Data Fig. 4d,e). As these embeddings are rich in biological and chemical information, initializing graph features with them is expected to improve performance in the challenging task of binding affinity prediction. When training on PDBbind, the GEMS baseline model, which lacks any language model embeddings, performed best on the CASF2016 test dataset. Incorporating language model features resulted in a continuous increase in cross-validation performance, without a corresponding improvement in test set performance. Conversely, when trained on PDBbind CleanSplit, the baseline model without language model features showed relatively low cross-validation and test dataset performance. However, the introduction of such features led to simultaneous improvements in both metrics. This suggests that PDBbind CleanSplit provides a better foundation for training protein-ligand affinity prediction models, as it eliminates straightforward paths to high test performance, such as exploiting biases and data leakages. Instead, the diversity inherent in the filtered dataset and the absence of structural similarities requires models to approach the task by actually learning the factors driving high-affinity proteinligand interactions. These models benefit from increased model complexity and enriched feature sets, such as those incorporating language model embeddings.

## Discussion

The PDBbind dataset remains the largest resource for training protein-ligand binding affinity prediction models. However, the development of a generalizable affinity prediction model requires refining this dataset to address its significant training redundancies and data leakage into the commonly used CASF bechmark. By developing a structure-based filtering algorithm, we created PDBbind CleanSplit, a refined training dataset with minimized redundancy and strict separation from the CASF complexes. With PDBbind CleanSplit, models can no longer rely on training data memorization, as all complexes resembling any from the CASF benchmarks have been excluded from the training dataset. Additionally, the removal of redundancy ensures that models are trained on a much more diverse dataset, ultimately improving their generalization capabilities. In summary, PDBbind CleanSplit provides an improved foundation for training binding affinity prediction models, setting a new standard for robust training and reliable evaluation in this field.

The impact of using PDBbind CleanSplit for training becomes evident in the performance drops of Pafnucy and GenScore, revealing that the true generalization capability of these previously top-performing model is much lower than reported. In contrast, our GEMS scoring function maintained excellent prediction accuracy when trained on PDBbind CleanSplit, achieving performance comparable to many deep-learning-based binding affinity prediction models that trained on the original PDBbind and profited from the associated data leakage. In addition, training GEMS was about 25 times faster than training Pafnucy and over 100 times faster than GenScore on the same GPU, thanks to our sparse graph-based modeling of protein-ligand interactions and an efficient GNN architecture. Combined with transfer learning from large language models, GEMS obtained an understanding of protein-ligand interactions and thus can better generalize to strictly external test datasets.

GEMS is a powerful scoring function designed to address a critical bottleneck in structure-based drug design (SBDD) and computational drug discovery. While recent generative models like AlphaFold3, RFdiffusion, and DiffSBDD can create libraries of novel protein-ligand interactions, their impact is constrained by the lack of tools to accurately predict binding affinities. Importantly, scoring *de novo* protein-ligand interactions demands models that go beyond exploiting structural similarities to existing complexes and demonstrate true generalization. GEMS fills this gap by offering robust scoring capabilities validated on strictly independent datasets, enabling identification of interactions with therapeutic potential.

We have prioritized accessibility by making our data, code, and model publicly available, including datasets of precomputed interactions graphs for fast reproduction of our results, and scripts for filtering the PDBbind database based on precomputed pairwise similarity matrices.

The current architecture and training regime of GEMS were specifically optimized to achieve state-of-the-art absolute binding affinity prediction on structurally diverse protein–ligand complexes in a data-leakage-free setting. For this reason, GEMS was trained on a dataset of high-quality structures with reduced redundancy, and it has not been exposed to docking poses or decoy conformations during training. As a result of this design, we recommend using classical scoring functions or machine learning models that have been trained on appropriately augmented datasets for tasks requiring fine discrimination between closely related docking poses, such as selecting the best conformation from many Glide or AutoDock Vina outputs. One avenue to improve GEMS in this direction is to augment its training data with high-quality docking poses and explicitly train the model to distinguish and rank closely related binding conformations. This would enhance the applicability of GEMS to docking-based virtual screening scenarios, where pose selection among many near-native docking conformations is usually required. Other desirable improvements include the ability to provide atom-level importance scores or optimize molecular conformations directly [53].

Promising future directions to further finetune PDBbind CleanSplit include the incorporation of more sophisticated methods for ligand similarity computation. While the current alignment and RMSE-based strategy employed in the filtering algorithm is effective at reducing data leakage and efficient enough to compute the 189 *×* 10^6^ pairwise similarities in PDBbind, its precision could be further improved using more advanced ligand comparison techniques, such as ROCS similarity scores [54].

## Methods

### Datasets

The main data resource used in this work was the PDBbind (v.2020) database, containing 19’443 protein-ligand complexes from the Protein Data Bank (PDB) with experimentally measured binding affinities. This database is split into a general set (n=14127) and a refined set (n=5316) that has been compiled based on strict curation criteria, including crystallographic structures (excluding NMR structures) with a resolution of *<* 2.5Å and an inhibition constant (Ki) or dissociation constant (Kd) in the range of 1pM to 10mM (pK range 2-12). To keep our training set as large as possible, we used a merged dataset containing all data from the general and the refined set as training data, excluding all complexes present in the Comparative Assessment of Scoring Functions (CASF) benchmark datasets (versions 2013 and 2016), which served as external test datasets in this work. Each complex is labeled with either an inhibition constant Ki, the dissociation constant Kd, or the half-maximal inhibitory concentration IC50. Despite concerns about the comparability of IC50 with Ki/Kd measurements [55], these metrics were considered interchangeable in this study and converted to pK values with *−* log_10_(*Ki/Kd/IC*50) to generate the final affinity labels. This decision is based on the observation that including IC50 complexes (n=6400) in the training data, despite being more noisy, has a small positive effect on the prediction performance of models on an independent test dataset (**Extended Data** Fig. 2)., confirming the findings of previous research [24].

During data preprocessing and graph construction, some protein-ligand complexes were excluded from the datasets based on the following criteria:

- Affinity label is not exact, e.g. *K_i_ <* 100*nm* (n=383)
- Error in RDKit when handling explicit hydrogens in some SDF files (n=45)
- Protein contains unknown residue or heteroatom in binding pocket, such as UNK or DOD (n=14)
- Error occurring during RDKit parsing due to incorrect valences in SDF files (n=5)
- Protein structure is not completely resolved and atoms are missing from the binding pocket (n=2)
- The ligand contains fewer than 5 heavy atoms (n=1)

This filtering reduced the size of the training dataset by 450 complexes to *N* = 18623 protein-ligand complexes. The CASF2016 (N=285) and CASF2013 (N=195) test sets were unaffected.

### Filtering Algorithm

To identify and remove structural similarities between the PDBbind database and the CASF benchmark datasets, our dataset filtering process relied on a combination of Tanimoto ligand similarity, protein similarity (TM scores), and a pocket-aligned ligand root-mean-square-deviation to compare the positioning of the ligands within the protein pockets. Tanimoto similarity is commonly used in cheminformatics to measure the similarity between small molecules. Based on comparing the chemical fingerprints, this score ranges between 0 (no similarity) and 1 (identical) and is a useful metric to identify compounds with similar structural and chemical properties. This score served as the first layer of our filtering, identifying pairs of complexes with similar ligands. The second layer of the filtering process used TM-align [43], a computational tool designed to compare protein structures by finding the optimal alignment of their three-dimensional shapes. It uses a scoring function based on the root-mean-square-distance of aligned residues and a length normalization factor, making it particularly effective for identifying structural similarities between proteins, regardless of sequence similarity. In our application, TM-align was valuable for identifying proteins that share similar binding pockets despite having low sequence identity. For instance, the test complex 1P1N and the training complex 3U92 share 53% sequence identity. Nevertheless, TM-align has recognized that 1P1N is well-represented within 3U92 and returns a TM score of 0.93. Alignment of the proteins then revealed that these complexes share identical binding pockets (Supplementary Fig. 1). The capability of our filtering process to identify complexes with similar interaction patterns in proteins that are otherwise dissimilar is a key advantage of our method over traditional sequence-based clustering approaches, which would overlook this similarity.

However, the combination of TM-score and Tanimoto similarity does not conclusively determine the similarity of the interaction patterns of two complexes. Even with a Tanimoto similarity of 1 and a TM-score of 1, two complexes might have different positioning of the ligand within the binding pocket, or the ligands might even bind at entirely different sites of the protein. Therefore, the third layer of our filtering process compares ligand positioning through a protein alignment followed by a root-mean-square-deviation (RMSD) calculation between the ligand atoms. For this analysis, one ligand’s atom coordinates are translated and rotated using the rotation matrix and translation vector obtained from TM-align (Supplementary Fig. 2). This alignment positions both ligands within the coordinate system of the optimal alignment of the protein structures. RMSD between the ligand atoms is then calculated to provide a quantitative measure of the positional similarity.

Our filtering algorithm imposes stringent rules on the structural and chemical similarity within the dataset. The similarity between two complexes is quantified on the basis of four computed similarity metrics:

- **Affinity:** The absolute difference between the reported binding affinities (pK values)
- **Tanimoto:** The Tanimoto similarity score to compare ligand structure was computed using RDKit library v.2024.03.3 based on count-based molecular fingerprints (GetCountFingerprint) of size 2048 and radius 2
- **TM-score:** TM-align [43] was used to assess protein structural similarity and to align complexes based on their protein residues. This algorithm identifies the best structural alignment between protein pairs and outputs a TM-score ranges from 0 (dissimilar structures) to 1 (identical structures) along with the translation vector and rotation matrix necessary to achieve optimal alignment. To find if one of the proteins is well represented within the other, we considered TM-scores normalized by the lengths of both amino acid chains and used the highest result as a similarity score.
- **RMSD:** A root-mean-square-deviation (RMSD) is computed to compare ligand positioning in the binding pocket of the proteins. For this, the protein structures were aligned together with the bound ligands by applying the translation vector and rotation matrix that TM-align used to generate an optimal protein alignment. Then, the atom coordinates of the aligned ligands were compared by computing the RMSD between the nearest points in the two point clouds. In case of different atom counts, the distances from each atom in the larger ligand to its nearest neighbor in the smaller ligand, resulting in larger RMSD values for ligands with different atom counts. A lower distance value indicates a very similar positioning of the two ligands in their binding pockets, and a higher value suggests very dissimilar binding conformations (or even binding at a different location of the protein).

#### Removal of Train-Test-Overlap

To remove the overlap between the training dataset (PDBbind) and the CASF test datasets (CASF2013 and CASF2016), our filtering algorithm removed all training complexes that are similar to any test complex in terms of protein structure, ligand structure, ligand binding and affinity. This eliminates data leakage by removing shared similarities between training and test datasets, effectively bringing the CASF datasets closer to being independent test datasets. A training complex was excluded if it shared all of the following similarities with a test complex:

1. The proteins have a TM-score higher than 0.8
2. The sum of the Tanimoto score (T) and the inverted RMSD is higher than 0.8 (*T* + (1 *− RMSD*))
3. The affinity labels are similar (*±*1 in pK units)

The first criterion ensures that training complexes are only excluded if they share a high structural similarity with a test complex. The second criterion balances ligand chemical similarity with binding conformation similarity. This approach means that a greater ligand positioning similarity is acceptable for two ligands with modest chemical similarity (e.g., Tanimoto coefficient, T=0.6). Conversely, for very similar ligands (T approaching 1), the training complex can be excluded from the dataset despite larger deviations in positioning. The third criterion ensures that training complexes are only excluded if they share a similar affinity label with the test complex. In addition, training complexes were excluded if they had an identical ligand (Tanimoto *>* 0.9) and a closely matching affinity label (*±*1) compared to a test complex. Together, this filtering excluded 711 training complexes. An overview over the model performance of GEMS, Pafnucy and the complex-based search algorithm in dependence of different dataset filtering thresholds for protein similarity and ligand similarity can be found in Extended Data Fig. 3.

#### Removal of Training Dataset Redundancy

A second filtering layer was applied to the training dataset to eliminate excessive redundancies while preserving the largest possible dataset size and quality. Initially, pairwise similarities between all training complexes were calculated using the described similarity metrics and recorded in a pairwise similarity matrix. To identify clusters of high similarity, this matrix was transformed into an adjacency matrix by applying the following thresholds:

1. The proteins have a TM-score higher than 0.8
2. The sum of the Taminoto score (T) and the inverted RMSD is higher than 1.3 (*T* + (1 *− RMSD*))
3. The affinity labels differ by less than *±*0.5 in pK unit

The resulting adjacency matrix connects all data points that meet these criteria. The filtering algorithm then iteratively removed complexes from the training dataset until no connections remained in the adjacency matrix. During each iteration, the complex with the highest number of connections (indicating the most similarities to other complexes) was excluded. In cases where multiple complexes had the same number of connections, preference was given to removing those from the PDBbind general set rather than the refined set. If ties persisted, the complex with the lowest resolution was preferentially excluded. In total, this filtering process excluded 1451 training complexes and allowed us to confidently train models using 5-fold cross-validation with random splitting, without the risk of inflated validation performance that could arise from similarities between training and validation datasets.

### Graph Construction

Each complex in the PDBbind database was translated into an affinity-labeled graph that models the interaction between the ligand and the protein (hereafter referred to as interaction graphs). For this, the affinity labels of all protein-ligand complexes in the database were extracted from the index files provided by the PDBbind database. The supplied inhibition constants (Ki), dissociation constant (Kd) and half-maximal inhibitory concentration (IC50) were converted to pK values with *−* log_10_(*Ki/Kd/IC*50) to generate the final affinity labels.

The interaction graphs were generated as follows: Protein PDB files were processed with Biopython v.1.83 to retrieve the amino acid sequences. Ligand SDF files were parsed with RDKit v.2024.03.3. From the atom coordinates of the ligand and protein, a pairwise distance matrix was generated to identify ligand atoms and protein atoms in close vicinity. An interaction distance of 5Å was considered sufficient to capture the most critical molecular interaction between ligand and protein. Consequently, if any atom of a protein residue was closer to a ligand atom than 5Å, the protein residue was considered to be in interaction distance to this ligand atom. Based on this, a list of amino acids and heteroatoms in interaction distance was generated for each ligand atom. To generate a graph representation of the protein-ligand interaction, a basic molecular graph was generated from the ligand by translating the atoms into graph nodes, and the covalent bonds into graph edges. To add an encoding of the protein pocket to these ligand graphs and model potential non-covalent interactions, all protein residues and heteroatoms previously determined to be in interaction distance were added as additional graph nodes and connected to all ligand atoms within interaction distance through additional edges. Importantly, the protein is represented at the amino acid level, which means that each amino acid is represented by a single graph node located at the position of its C*α* atom. The resulting interaction graphs consisted of an atom-level graph representation of the ligand, supplemented with an amino acid-level representation of its surrounding protein residues (Supplementary Fig. 3). All edges of these basic interaction graphs were undirected (applying message-passing in both directions), and self-loops were included for each node.

#### Featurization

The interaction graphs’ initial node and edge features consisted of one-hot-encoded chemical properties computed with the RDKit v.2024.03.3 library. The following basic chemical features were included:

- Nodes: Atom type (B, C, N, O, P, S, Se, metal, halogen), ring membership, hybridization, formal charge, aromaticity, atomic mass, number of bonded hydrogens, degree, chirality
- Edges: Edge type (covalent, self-loop, non-covalent), length, bond type (single/double/triple/aromatic), conjugation, ring membership, stereochemistry

In feature vectors of edges connecting a ligand atom node with a protein residue node, the length feature was replaced with four values representing the distances between the ligand atom and the four main backbone atoms of the residue (N, C*α*, C and virtual C*β*). If any feature did not apply to a certain node (e.g. atom type for a node representing a protein residue), the feature was replaced with zero padding to maintain consistent feature dimensions across the graphs.

The features of the nodes representing protein residues were additionally supplemented with a vector containing the one-hot-encoded amino acid type and with amino acid embeddings. These embeddings were generated with ESM2 (T6 8M checkpoint downloaded from huggingface https://huggingface.co/facebook through the transfomers library v.4.33.3) [50] and ANKH (base model downloaded through the ankh python library v.1.10.0) [51]. For this, the amino acid sequence of a protein was passed through the downloaded tokenizers and model checkpoints. The resulting matrices contained embeddings for each amino acid in the protein sequences, which were then appended to the features of the corresponding graph nodes.

In addition, a ligand embedding was computed for each complex using the ChemBERTa-2 language model (ChemBERTa-77M-MLM downloaded from https://huggingface.co/DeepChem through the transformers library v.4.33.3) [52]). The smiles codes of all ligands in the dataset were generated with RDKit v.2024.03.3 and passed through the downloaded model to obtain ligand embeddings. These embeddings were used to initialize the graph’s initial global feature.

### Model Architecture

Our graph neural network models were implemented using PyTorch v.2.0.1 and Torch Geometric (pyg) v.2.5.2. The model architecture captures and integrates multi-level graph information across nodes, edges, and the entire graph. It receives batches of interaction graphs as input and alternates between updating the graph’s edge features, node features, and global features (**Fig. 4b**). The architecture includes the following components:

1. A NodeTransformMLP module, which applies a multi-layer perceptron (MLP) with ReLU and dropout to transform the input node features.
2. An EdgeModel module to update edge features. It processes the concatenated features of source nodes, destination nodes, and existing edge features through an MLP with ReLU and dropout.
3. A NodeModel module to update node features. It uses a graph attention network (GATv2Conv) [56] convolution to update node features based on their neighboring nodes and edge attributes.
4. A GlobalModel module concatenates the global graph features with aggregated node features and applies an MLP with dropout to create updated global graph features.

The entire model architecture of GEMS encompasses five convolutional layers, of which three update the node features and two update the edge features, resulting in a model architecture with 1’032’129 learnable parameters. In a forward pass of the model, the node features are first transformed by the NodeTransformMLP module. These transformed node features, along with edge attributes and global features, are then processed through two consecutive graph layers. Each graph layer performs the following sequence of updates: the EdgeModel updates the edge features based on the connected node’s features, the NodeModel updates the node features based on the features of neighboring nodes and the connecting edges, and the GlobalModel updates the global features with pooled node features. After the first graph layer, batch normalization is applied to the node, edge, and global features. Following the second graph layer, a dropout layer is applied to the global features to prevent overfitting. Finally, the global features are passed through two fully connected layers with a ReLU activation to produce the final output.

As graph convolutional operator implemented in the NodeModel module, GATv2Conv [56] from Torch Geometric was selected with concatenation of multi-head attention. This convolutional layer computes updated node features *x*^′^*_i_* for node *i* with

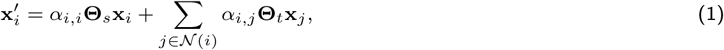

where **x***_j_* represents the feature vector of the neighboring node *j*, *N* (*i*) the set of neighboring nodes of node *i*, **Θ***_s_* a learnable weight matrix applied to the features of the source node *i*, **Θ***t* a learnable weight matrix applied to the features of the target node *j* and *α_i,j_* the attention coefficients representing the importance of node *j*’s features to node *i*. The attention coefficients *α_i,j_* are computed as

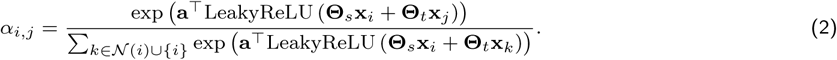

where LeakyReLU is an activation function applied to introduce non-linearity. Taken together, this layer updates each node’s features by aggregating the transformed features of its neighbors and itself, weighted by attention coefficients that are dynamically computed based on the features of both the source and target nodes. This mechanism allows the model to focus on the most relevant parts of the graph structure during learning.

To find the optimal number of message-passing steps, channels and attention heads of the individual layers, different model architectures were tested:

- Number of message passing steps: Search space [1,2,3] with final value 2.
- Graph pooling operator: Search space [mean pooling, add pooling, max pooling] with final choice add pooling.
- NodeTransformMLP output channels: Search space [256, 128, 64] with final value 64.
- NodeModel output channels: Search space [256, 128, 64] with final value 64.
- EdgeModel output channels: Search space [128, 64] with final value 64.
- GlobalModel output channels: Search space [768, 512, 384, 256] with final value 384.

### Model Training and Selection

During the training of GEMS and Pafnucy, all model variants trained on the same dataset were subjected to the same five-fold cross-validation split to eliminate the variability introduced by differences in data partitioning. The models were trained across all five splits, and the models that achieved the lowest validation root-mean-square-error (RMSE) were saved. The training objective was to minimize the RMSE of the predicted pK values of the training complexes, using a stochastic gradient descent (SGD) optimizer. To prevent overfitting, early stopping was implemented. This technique halted the training process if there was no improvement in the validation RMSE for 100 consecutive epochs. All models were trained on one NVIDIA GeForce RTX 4090 or one NVIDIA GeForce RTX 3090 for approximately 200-1200 epochs (depending on the early stopping), taking between 10 and 60 minutes. The training hyperparameters were optimized as follows:

- Batch size: Search space [128,256,512,640] with final value 256.
- Learning rate: Search space [0.0001, 0.001, 0.01] with final value 0.001.
- Weight decay: Search space [0.0001, 0.001, 0.01] with final value 0.001.

From all trained models, the one with the highest and most consistent validation performance across all five folds was selected. This ensures that the model with the most robust generalization across different subsets of our training data is selected. For testing models on the CASF test datasets, the CASF complexes were passed through all five cross-validation models and the predictions from all five models were averaged to generate the final ensemble predictions.

To robustly determine the uncertainty of GEMS, we trained it using our five-fold cross-validation approach at five different random seeds. This approach ensures that the randomness in the data splitting differs with each training run, which allows us to estimate the variability in performance that arises from different data splits. By averaging the outcomes and calculating the standard deviation between these iterations, we generated error bars for our performance metrics.

#### Ablation

To facilitate ablation experiments, our script for constructing interaction graph datasets from protein-ligand complexes includes a parameter that allows for the optional removal of all protein information from the graphs. To test whether GEMS relies on the presence of both ligand and protein data, we generated ligand-only versions of the PDBbind, PDBbind CleanSplit, and CASF datasets by removing all protein nodes, leaving only the molecular graph of the ligand. We then trained and tested GEMS as described above using these ligand-only datasets.

#### Pafnucy

The Pafnucy model was obtained from the authors public repository (https://gitlab.com/cheminfIBB/pafnucy) and trained on a NVIDIA GeForce RTX 3090 GPU using the provided scripts and instructions. Protein-ligand complexes were preprocessed and datasets prepared according to the authors’ specifications. We evaluated its performance on the training and validation sets, as well as on the CASF2016 dataset, using the predictions from the output text file generated by the training script. To compare the performance of our model with that of Pafnucy, we used the same cross-validation method and identical data split.

#### GenScore

The GenScore model was obtained from the authors public repository (https://github.com/sc8668/GenScore) and preprocessed graph datasets covering the full PDBbind database (version 2020) and the CASF2016 benchmark were downloaded from Zenodo (https://zenodo.org/records/7578480). Using these datasets, we trained and evaluated GenScore by running the provided training and inference scripts with default settings on a NVIDIA GeForce RTX 3090 GPU. To assess GenScore’s scoring power on CASF2016, we employed a benchmarking script from the official CASF2016 scoring power evaluation protocol [10]. Although GenScore produces a numerical scores that are not on the same scale as experimental binding affinities (pK values), the benchmarking script applies a simple linear regression to map the predicted scores to the experimental pK scale. From this regression model, we obtained the predicted binding affinity values for each complex, which allowed us to compute absolute prediction errors and RMSE values, as shown in **Fig. 3**. Since the provided GenScore code supports only random train-validation splits, we could not reproduce the exact data splits used in the GEMS and Pafnucy experiments. However, to enable the fairest possible comparison under these constraints, we trained five GenScore models using different random seeds and reported the mean performance and standard deviation across these runs. To train GenScore on the filtered PDBbind CleanSplit dataset, we filtered the provided training dataset to contain only the complexes in CleanSplit and repeated the above procedure.

## Data availability

PDBbind data are available at: http://www.pdbbind.org.cn/. The data generated in this study are freely available on GitHub (https://github.com/camlab-ethz/GEMS) including GEMS model parameters and PDBbind CleanSplit. For fast reproduction of our results, we provide PyTorch datasets of precomputed interaction graphs for the entire PDBbind database on Zenodo (https://doi.org/10.5281/zenodo.14260170). To enable quick establishment of leakage-free evaluation setups with PDBbind, we also provide pairwise similarity matrices for the entire PDBbind dataset on Zenodo.

## Code availability

The code generated in this study is freely available on GitHub. All scripts for dataset filtering, model training and inference can be accessed through https://github.com/camlab-ethz/GEMS. A docker container for easy reproduction and implementation is provided.

## Acknowledgements

This study was financed by C.H.’s ZHAW DIZH Fellowship (Call 2022) and supported by NCCR Catalysis, a National Centre of Competence in Research funded by the Swiss National Science Foundation (Grant No. 225147 to R.B. und P.S). S.M.’s contribution to this work was supported in part by the DOE SEA-CROGS project (DE-SC-0023191).

## Author contributions

D.G., C.H. and S.M. conceived the GEMS model architecture. D.G., P.S., F.M., C.H., S.M. and R.M.B designed the experiments. D.G. carried out the experiments and D.G., P.S., R.M.B. and S.M. analysed the data. P.S. tested the code for usability with different datasets and set up a docker container. D.G., C.H., S.M. and R.M.B. wrote the manuscript with feedback from F.M. and P.S. The project was supervised by C.H., R.M.B. and S.M.

## Extended Data

**Extended Data Figure 1:**
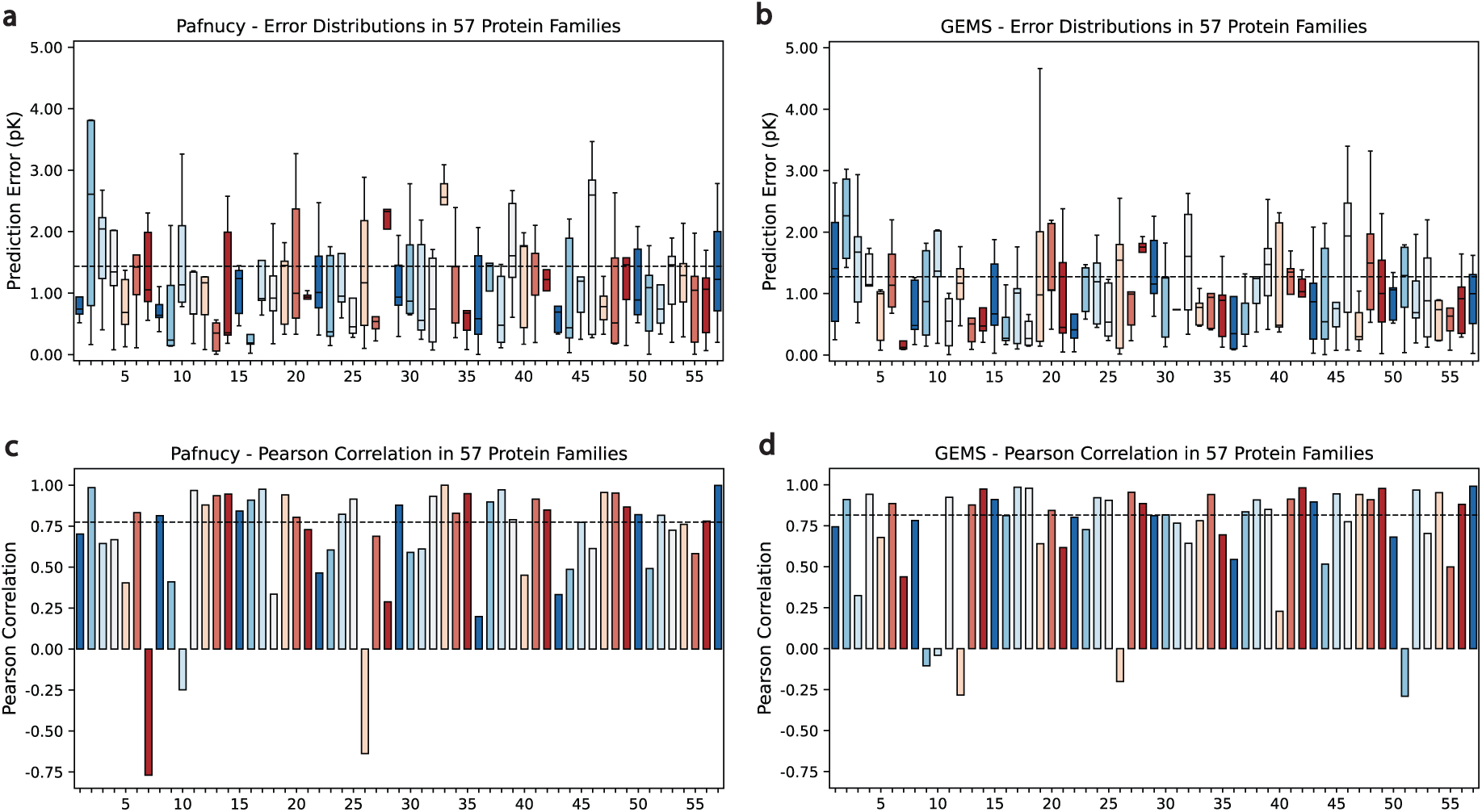
Performance of Pafnucy and GEMS in Different Protein Families: Comparison of the prediction accuracy of Pafnucy and GEMS per protein cluster in the CASF2016 benchmark dataset (N=285). **a)** and **b)** Distribution of absolute prediction errors (pK unit) of Pafnucy and GEMS within 57 protein clusters, where each cluster contains five proteins of the same family with ligands spanning a wide range of binding affinities. The black dashed lines indicate the prediction RMSE on the entire CASF2016 dataset. **c)** and **d)** Prediction Pearson correlations of Pafnucy and GEMS within the same 57 protein clusters. Black dashed lines indicate the Pearson correlation on the entire CASF dataset.

**Extended Data Figure 2:**
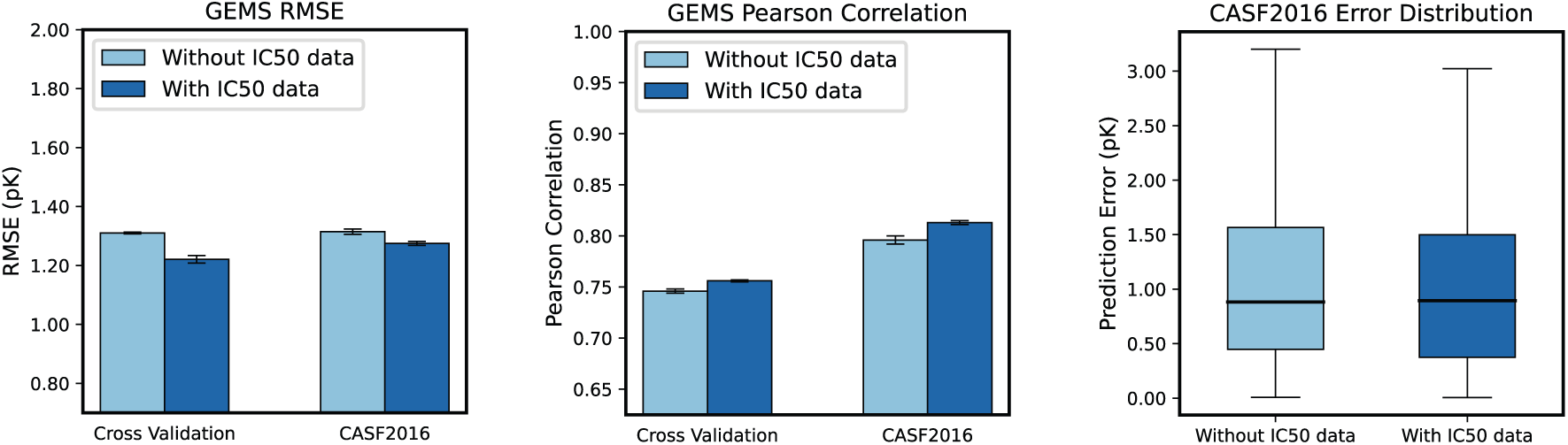
Performance of GEMS with and without IC50-labeled complexes: Comparison of cross validation and CASF2016 (N=285) prediction accuracy of GEMS in terms of **a)** RMSE, **b)** Pearson correlation and **c)** absolute prediction errors depending on inclusion of 6400 complexes labeled with IC50 values in the training data. Performance values represent the mean values of five models trained with 5-fold cross-validation at different random seeds. Error bars represent data uncertainty, calculated as the standard deviation of the performance across the five models

**Extended Data Figure 3:**
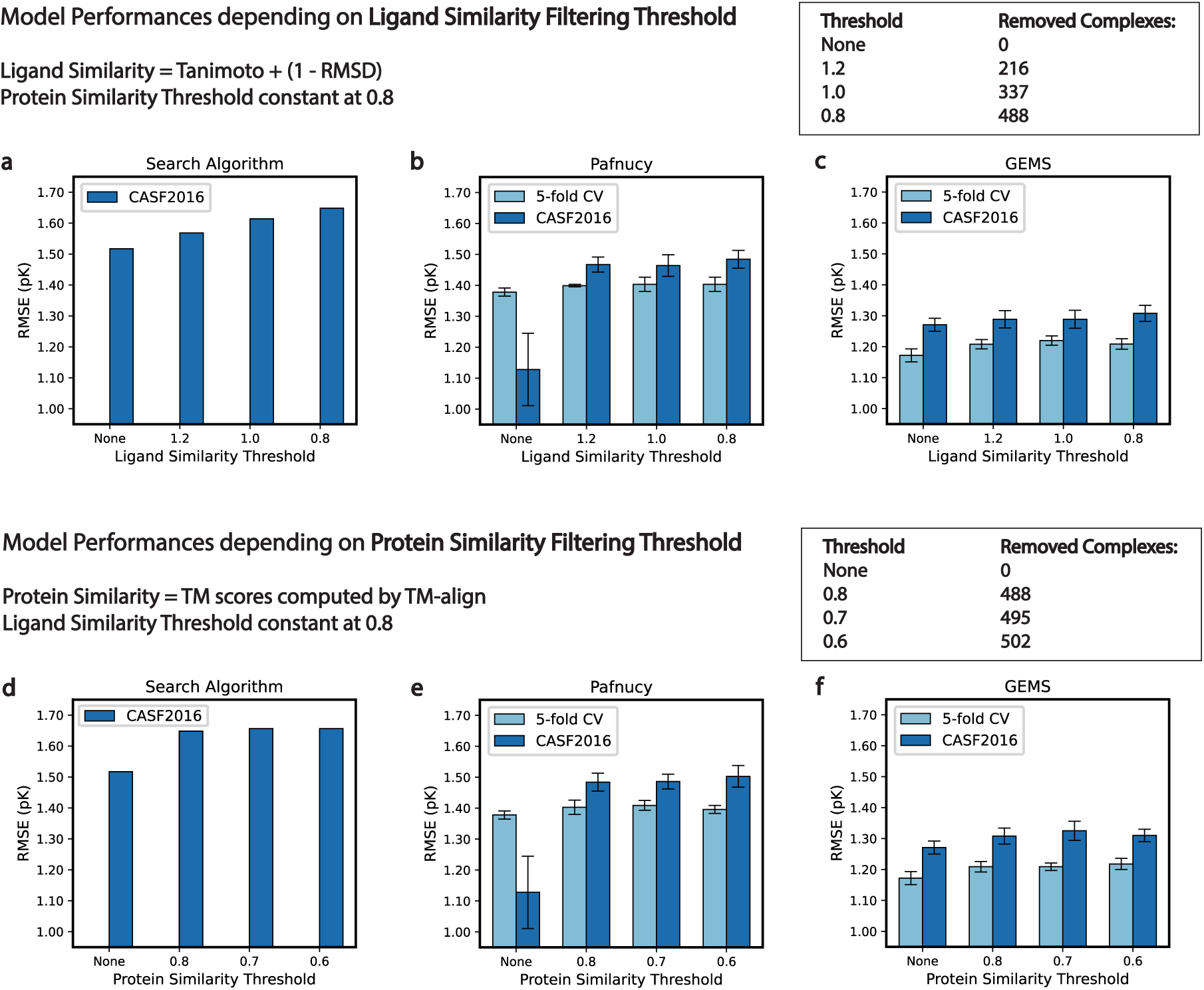
Influcence of Filtering Thresholds on Model Performance: Changes in cross-validation and CASF2016 (N=285) prediction accuracy of the complex-based search algorithm, Pafnucy and GEMS in dependence of different training dataset filtering thresholds for protein similarity and ligand similarity. **a), b) and c)**: Changes in model performance with varying ligand similarity thresholds at constant protein similarity threshold 0.8. **d), e) and f)** Changes in model performance with varying protein similarity thresholds at constant ligand similarity threshold 0.8. Ligand similarity reflects a combined assessment of Tanimoto scores and pocket-aligned ligand RMSD (Tanimoto + 1-RMSD). Protein similarity represents TM-scores computed by TM-align. The first group of bars in each plot represents the model performance without any training dataset filtering. The performance values in **b), c), e) and f)** reflect the mean validation and CASF2016 RMSE of five models originating from 5-fold cross-validation, with error bars representing the standard deviation across the performance of the five models. To ensure a fair comparison, Pafnucy and GEMS were trained using the exact same 5-fold cross-validation split.

**Extended Data Figure 4:**
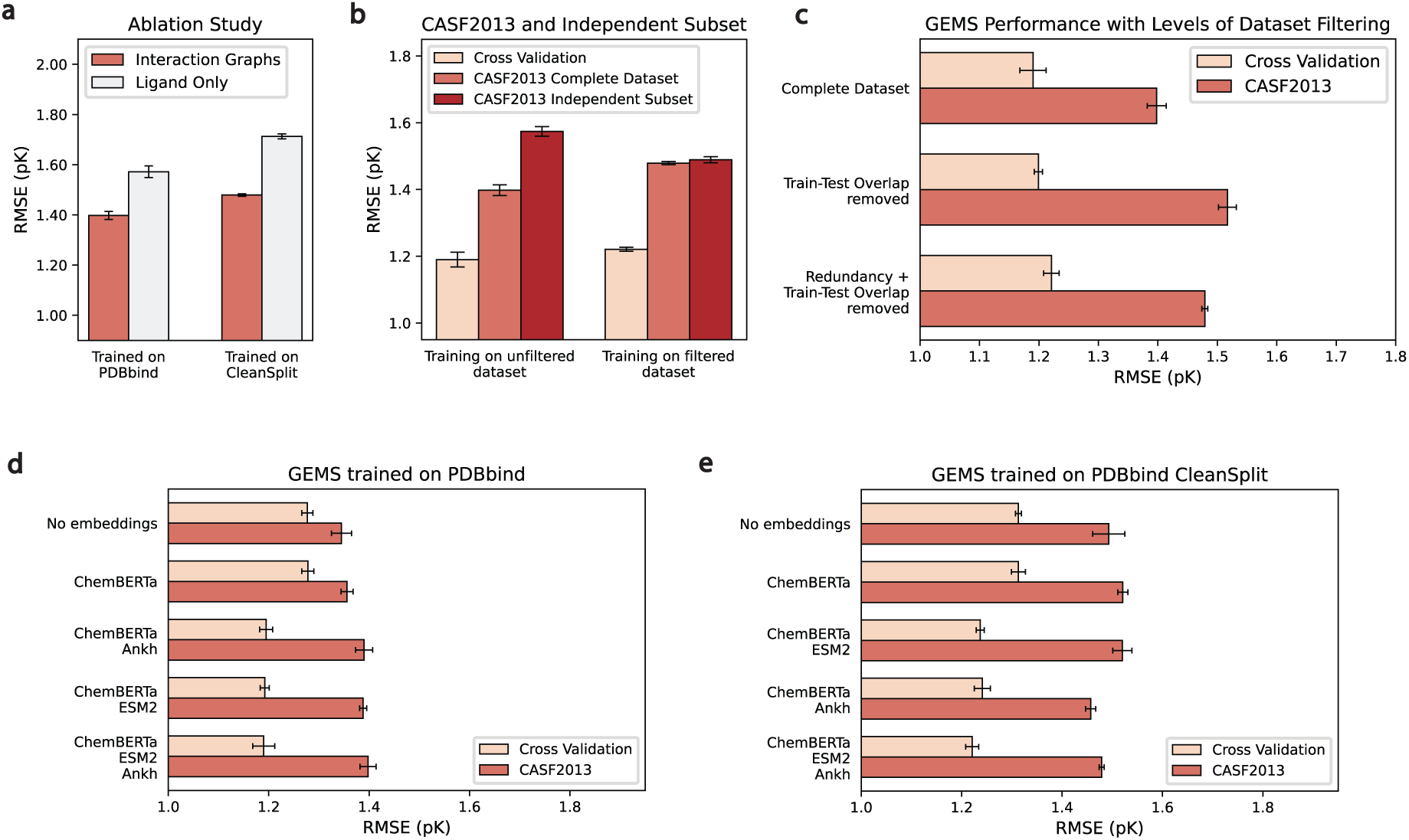
Impact of Ablation and Dataset Filtering on GEMS Performance on CASF2013 (N=195): **a)** Ablation study showing the impact of removing all protein information from the graphs on CASF2013 performance, comparing models trained on PDBbind (left) and PDBbind CleanSplit (right). **b)** Comparison of GEMS performance on cross-validation (CV), CASF2013, and the independent subset of CASF2013 (N=89), for models trained on PDBbind (left) and PDBbind CleanSplit (right). **c)** CASF2013 performance of GEMS with varying levels of training dataset filtering: complete dataset (PDBbind), train-test overlap removed, and both overlap and redundancy removed (PDBbind CleanSplit). Error bars represent data uncertainty, calculated as the standard deviation of the performance across five models trained with 5-fold cross-validation at different random seeds. **d)** Effect of incorporating language model embeddings on the cross-validation and CASF2013 (n=285) performance of GEMS models trained on the original PDBbind dataset. **e)** Effect of incorporating language model embeddings on the cross-validation and CASF2013 (n=285) performance of GEMS models trained on the PDBbind CleanSplit.

