## Supplemental Information for "GEMS – Enhancing Generalizable Binding Affinity Prediction by Removing Data Leakage and Integrating Language Model Embeddings into Graph Neural Networks"

### Supplementary Information

#### Supplementary Figure 1 - Detection of Similar Interaction Patterns Despite Low Sequence Identity

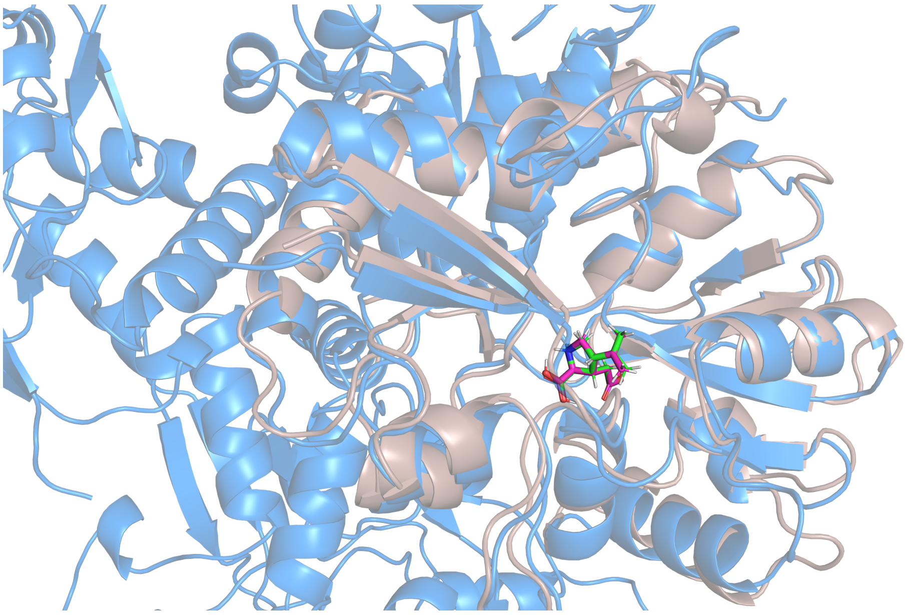

**Supplementary Figure 1: Detection of Similar Interaction Patterns Despite Low Sequence Identity:** Superposition of the test complex 1P1N (protein in gray, ligand in magenta) with the training complex 3U92 (protein in blue, ligand in green) structurally aligned with TM-align. These complexes have a low sequence identity of 53% but a high TM-score of 0.93, as 1P1N is a substructure of 3U92. They contain closely matching binding pockets, identical ligands, and similar binding conformations. Due to this substantial similarity, these complexes would provide nearly identical input data points to our models. Considering the comparable affinity labels, complex 3U92 was excluded from the training dataset to eliminate train-test data leakage. The capability to identify complexes with similar interaction patterns despite low sequence identity is a key advantage of our filtering algorithm over traditional sequence-based methods.

#### Supplementary Figure 2 - Pocket-Aligned Ligand RMSD Calculation

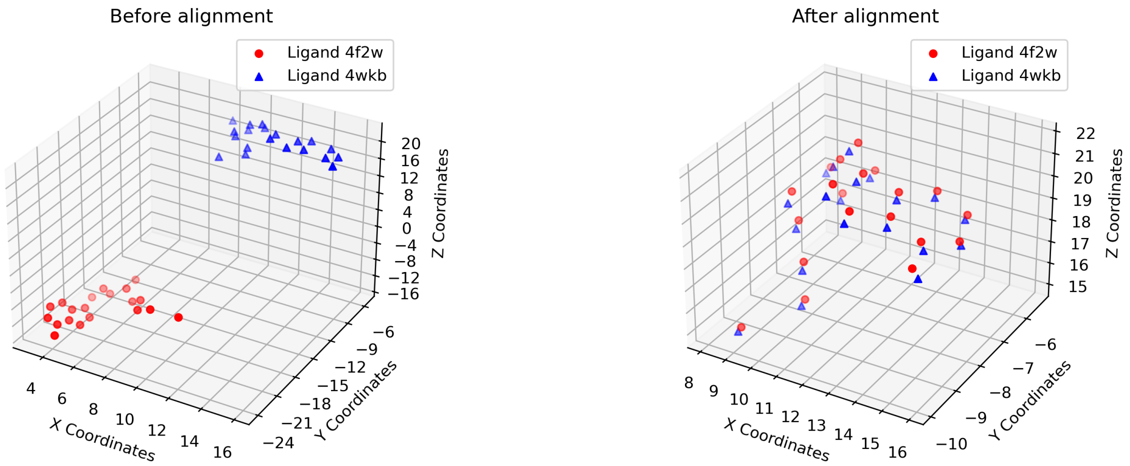


**Supplementary Figure 2: Ligand Binding Conformation Similarity with Pocket-Aligned RMSD:** a) Atom coordinates of the ligands of 4F2W and 4WKB before protein alignment. b) Atom coordinates of the ligands of 4F2W and 4WKB after protein alignment. The complexes 4F2W (CASF2016) and 4WKB (PDBbind) are highly similar, with a pK difference of 0.27, a Tanimoto score of 1.0 and a TM-score of 0.99, indicating identical ligands and nearly identical proteins. However, to conclusively assess the structural similarity of these complexes, it is necessary to compare the binding conformations of the ligands. For this, the complex 4F2W is translated and rotated into the coordinate system of the optimal protein alignment using the translation vector and rotation matrix returned by TM-align. This protein-alignment also aligns the contained ligands and reveals their nearly identical binding conformations (RMSD=0.33Å).

### Supplementary Figure 3 - Interaction Graphs

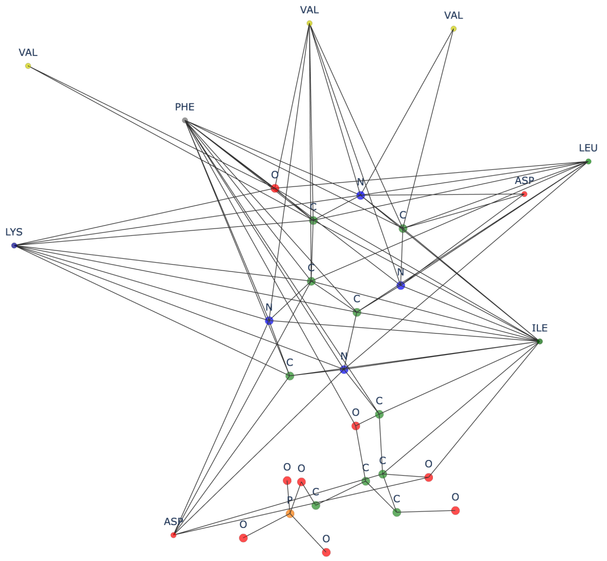


**Supplementary Figure 3: Interaction Graph Generated From the 5kam Protein-Ligand-Complex** This example graph includes a graph representation of the ligand molecules, where atoms are represented as nodes and bonds as edges, and a residue-level representation of the protein pocket. Edges between ligand atoms represent covalent bonds, while edges connecting ligand atoms with amino acids denote spatial proximity, suggesting potential non-covalent interactions between them.

### Supplementary Note - Training Behaviour Changes with Dataset Filtering

During the training of GEMS, all model variants were subjected to the same five-fold cross-validation procedure. Among all tested variants of GEMS, the variant with the highest and most consistent validation performance across all five folds was selected for testing on the CASF test dataset. Using the cross-validation performance as a selection criteria ensures the choice of the model showing the most robust generalization across different subsets of our training data, making it the most likely to generalize to new data. However, for GEMS models trained on the original PDBbind, the cross-validation performances did not correlate positively with the test set outcomes. Many models with moderate or low cross-validation performance achieved top-tier results on CASF2016 (RMSE of up to 1.15) and CASF2013 (RMSE up to 1.265), which is, to our knowledge, the best performance on CASF2013 reported to date. Despite these excellent benchmark metrics, we disregarded these models and focused on the models with highest cross-validation performance, as these are most likely to generalize successfully to new data.
This discrepancy between cross-validation results and actual test performance is concerning, as 5-fold cross-validation is generally considered a strong indicator of generalization. Notably, these high-performing models achieved their best test results with minimal dropout, whereas increasing dropout improved cross-validation but reduced test performance. This suggests that these models trained on PDBbind rely on overfitting and memorizing training data. Due to the train-test overlap, overfitting to the training data also effectively boosts test dataset performance, resulting in models with exceptional benchmark performance.

In contrast to the models trained on PDBbind, the GEMS models trained on PDBbind CleanSplit showed a closer correlation between 5-fold cross-validation performances and CASF test performance. The models that achieved the highest cross-validation performance consistently showed the best test set results. Additionally, these models typically achieved the highest validation and test performances with higher levels of dropout, highlighting that the prevention of overfitting is crucial for enhancing their test performance. This indicates that GEMS models trained on PDBbind CleanSplit do not rely on memorization, but rather on an understanding of the factors that contribute to high-affinity protein-ligand interactions.

### Supplementary Note - Graph Construction and Featurization

To train a robust prediction model on structural protein-ligand data, a sparse, rotation and translation-invariant encoding of the structural data is vital to allow models to learn on this data in a parameter-efficient way. We used graph representations to model the interaction in a protein-ligand complex. The core of these graph representations is an atom-level molecular graph of the ligand molecule, which is extended with an amino acid-level graph representation of the protein pocket. The exclusion of protein residues not involved in ligand binding and the sparse modeling of the protein pocket on amino acid level significantly reduced the size and complexity of the molecular graphs while preserving essential interaction data. In addition, the reduced complexity of the graphs lead to very fast model training compared to 3D-CNNs and GCN models including more detailed atom-level protein representations. The performance of GEMS on strictly separated test datasets indicates that modeling protein-ligand interactions at this level of detail is sufficient to make accurate predictions of binding affinity.

Nevertheless, the main advantage of representing amino acids as single nodes is the possibility to featurize these nodes with amino acid embeddings derived from protein language models. These models have been trained on vast collections of protein data, making the generated embeddings rich in biological and structural information. We hypothesized that incorporating these amino acid embeddings would significantly increase the predictive power of our models. Indeed, our results demonstrate that models trained on graphs featurized with language model embeddings significantly outperform baseline models that lack these features (see **Figure 5c**). By leveraging these embeddings, we can greatly enhance the feature set for our machine learning tasks, leading to more accurate predictions of protein-ligand binding affinities.

### Supplementary Note - Model Architecture

In our research, we employ a graph convolutional network (GCN) architecture designed to efficiently and effectively process graph-structured molecular data. The core model is a graph attention (GAT) network combined with multi-layer perceptrons (MLP) for updating edge features. Initially, node and edge features undergo transformation via MLPs, followed by a sequence of alternating updates for nodes and edges. After each edge feature update, the node features are updated using graph attention network (GATv2Conv) convolution, followed by an update of the global features. This sequence of operations allows our model to capture multi-level information from individual nodes, edges, and the entire graph structure. The global graph features are dynamically updated throughout the process, integrating node representations based on different neighborhood ranges. This combination of detailed local information with broader global information allows the model to integrate both local and global connectivity patterns into the final graph representation. These architectural features make this model setup particularly suitable for molecular graph-level tasks such as protein-ligand binding affinity prediction.

### Supplementary Table - Clusters in CASF2016

| ID | PDB codes in this cluster with Binding Constant ($logK_a$) | Protein |
|---|---|---|
| 1 | 2XB8 (7.59), 3N76 (6.85), 3N7A (3.70), 3N86 (5.64), 4CIW (4.82) | 3-Dehydroquinate dehydratase |
| 2 | 1NC1 (6.12), 1NC3 (5.00), 1Y6R (10.11), 4F2W (11.30), 4F3C (11.82) | 5'-Methylthioadenosine/s-adenosylhomocysteine nucleosidase |
| 3 | 3U8K (8.66), 3U8N (10.17), 3ZDG (7.10), 4QAC (9.40), 3WTJ (6.53) | Acetylcholine-binding protein |
| 4 | 1E66 (9.89), 1GPK (5.37), 1GPN (6.48), 1H22 (9.10), 1H23 (8.35) | Acetylcholinesterase |
| 5 | 2WVT (6.12), 2XII (7.20), 4J28 (5.70), 4JFS (5.27), 4PCS (7.85) | Alpha-L-fucosidase |
| 6 | 1PS3 (2.28), 3D4Z (4.89), 3DX1 (3.58), 3DX2 (6.82), 3EJR (8.57) | Alpha-mannosidase 2 |
| 7 | 1Z95 (7.12), 3B5R (8.77), 3B65 (9.27), 3B68 (8.40), 3G0W (9.52) | Androgen receptor |
| 8 | 3QQS (5.82), 3R88 (4.82), 3TWP (3.92), 4GKM (5.17), 4OWM (2.96) | Anthranilate phosphoribosyltransferase |
| 9 | 2FXS (6.06), 2IWX (6.68), 2VW5 (8.52), 2WER (7.05), 2YGE (5.06) | Atp-dependent molecular chaperone hsp82 |
| 10 | 2CBV (5.48), 2CET (8.02), 2J78 (6.42), 2J7H (7.19), 2WBG (4.45) | Beta-glucosidase a |
| 11 | 2R9W (5.10), 3GR2 (2.52), 3GV9 (2.12), 4JXS (4.74), 4KZ6 (3.10) | Beta-lactamase |
| 12 | 3G2Z (2.36), 3G31 (2.89), 4DE1 (5.96), 4DE2 (4.12), 4DE3 (5.52) | Beta-lactamase CTX-M-9a |
| 13 | 3NQ9 (4.03), 3UEU (5.24), 3UEV (5.89), 3UEW (6.31), 3UEX (6.92) | Beta-lactoglobulin |
| 14 | 2VKM (8.74), 3RSX (4.41), 3UDH (2.85), 4DJV (6.72), 4GID (10.77) | Beta-secretase 1 |
| 15 | 1K1I (6.58), 1O3F (7.96), 1UTO (2.27), 3GY4 (5.10), 4ABG (3.57) | Beta-trypsin |
| 16 | 3P5O (7.30), 3U5J (5.61), 4LZS (4.80), 4OGJ (6.79), 4WIV (6.26) | Bromodomain-containing protein 4 |
| 17 | 3UI7 (9.00), 3UUO (7.96), 4LLX (2.89), 5C28 (5.66), 5C2H (11.09) | cAMP and cAMP-inhibited cGMP 3',5'-cyclic phosphodiesterase 10A |
| 18 | 1Q8T (4.76), 1Q8U (5.96), 1YDR (5.52), 1YDT (7.32), 3AG9 (8.05) | cAMP-dependent protein kinase catalytic subunit alpha |
| 19 | 2WEG (6.50), 3DD0 (9.00), 3KWA (4.08), 3RYJ (7.80), 4JSZ (2.30) | Carbonic anhydrase 2 |
| 20 | 3NW9 (9.00), 3OE4 (7.47), 3OE5 (6.88), 3OZS (5.33), 3OZT (4.13) | Catechol o-methyltransferase |
| 21 | 1PXN (7.15), 2FVD (8.52), 2XNB (6.83), 3PXF (4.43), 4EOR (6.30) | Cell division protein kinase 2 |
| 22 | 4AGN (3.97), 4AGP (4.69), 4AGQ (5.01), 5A7B (3.57), 5ABA (2.98) | Cellular tumor antigen p53 |
| 23 | 3ARP (7.15), 3ARQ (6.40), 3ARU (3.22), 3ARV (5.64), 3ARY (6.00) | Chitinase a |
| 24 | 1LPG (7.09), 1MQ6 (11.15), 1Z6E (9.72), 2XBV (8.43), 2Y5H (5.79) | Coagulation factor x heavy chain |
| 25 | 4CR9 (4.10), 4CRA (7.22), 4CRC (8.72), 4TY7 (9.52), 4X6P (8.30) | Coagulation factor xi |
| 26 | 2ZCQ (8.82), 2ZCR (6.87), 2ZY1 (7.40), 3ACW (4.76), 4EA2 (6.44) | Dehydrosqualene synthase |
| 27 | 2V00 (3.66), 3PRS (7.82), 3PWW (7.32), 3URI (9.00), 3WZ8 (5.82) | Endothiapepsin |
| 28 | 1QKT (9.04), 2P15 (10.30), 2POG (9.54), 2QE4 (7.96), 4MGD (4.69) | Estrogen receptor |
| 29 | 1P1N (6.80), 1P1Q (4.89), 1SYI (5.44), 2AL5 (8.40), 4U4S (2.92) | Glutamate receptor 2 |
| 30 | 1VSO (4.72), 3FV1 (9.30), 3FV2 (8.11), 3GBB (6.90), 4DLD (5.82) | Glutamate receptor, ionotropic kainate 1 |
| 31 | 3EBP (5.91), 3G2N (4.09), 3L7B (2.40), 3SYR (5.10), 4EKY (3.52) | Glycogen phosphorylase, muscle form |
| 32 | 1YC1 (6.17), 2XDL (3.10), 2YKI (9.46), 3B27 (5.16), 3RLR (7.52) | Heat shock protein hsp 90-alpha |
| 33 | 3AO4 (2.07), 3ZSO (5.12), 3ZSX (3.28), 3ZT2 (2.84), 4CIG (3.67) | Integrase |
| 34 | 3EHY (5.85), 3LKA (2.82), 3NX7 (8.10), 3TSK (7.17), 4GR0 (9.55) | Macrophage metalloelastase |
| 35 | 2ZB1 (6.32), 3E92 (8.00), 3E93 (8.85), 4DLI (5.62), 4F9W (6.94) | Mitogen-activated protein kinase 14 |
| 36 | 2VVN (7.30), 2W4X (4.85), 2W66 (4.05), 2WCA (5.60), 2XJ7 (6.66) | O-glcnacase $BT_4395$ |
| 37 | 3COY (6.02), 3COZ (5.57), 3IVG (4.30), 4DDH (3.32), 4DDK (2.29) | Pantothenate synthetase |
| 38 | 2P4Y (9.00), 2YFE (6.63), 3B1M (8.48), 3FUR (8.00), 3U9Q (4.38) | Peroxisome proliferator-activated receptor gamma |
| 39 | 1A30 (4.30), 1EBY (9.70), 2QNQ (6.11), 3O9I (11.82), 1G2K (7.96) | HIV-1 protease |
| 40 | 1R5Y (6.46), 1S38 (5.15), 3GC5 (7.26), 3GE7 (8.70), 3RR4 (4.55) | Queuine tRNA-ribosyltransferase |
| 41 | 1O0H (5.92), 1U1B (7.80), 1W4O (5.22), 3D6Q (3.76), 3DXG (2.40) | Ribonuclease, pancreatic |
| 42 | 3CJ4 (6.51), 3GNW (9.10), 4EO8 (8.15), 4IH5 (4.11), 4IH7 (5.24) | RNA-directed RNA polymerase |
| 43 | 2WTV (8.74), 3E5A (8.23), 3MYG (10.70), 3UO4 (6.52), 3UP2 (7.40) | Serine/threonine-protein kinase 6 |
| 44 | 1NVQ (8.25), 2BR1 (5.14), 2BRB (4.86), 3JVR (5.72), 3JVS (6.54) | Serine/threonine-protein kinase Chk1 |
| 45 | 2C3I (7.60), 3BGZ (6.26), 3JYA (6.89), 4K18 (8.96), 5DWR (11.22) | Serine/threonine-protein kinase pim-1 |
| 46 | 2WN9 (8.52), 2WNC (6.32), 2X00 (11.33), 2XYS (7.42), 2YMD (3.16) | Soluble acetylcholine receptor |
| 47 | 3KR8 (8.10), 4J21 (7.41), 4J3L (7.80), 4KZQ (6.10), 4KZU (6.50) | Tankyrase-2 |
| 48 | 1QF1 (7.32), 1Z9G (5.64), 3FCQ (2.77), 4TMN (10.17), 5TMN (8.04) | Thermolysin |
| 49 | 1BCU (3.28), 1OYT (7.24), 2ZDA (8.40), 3BV9 (5.36), 3UTU (10.92) | Thrombin light chain |
| 50 | 4BKT (3.62), 4W9C (4.65), 4W9H (6.73), 4W9I (5.96), 4W9L (5.02) | Transcription elongation factor b polypeptide 2 |
| 51 | 3F3A (4.19), 3F3C (6.02), 3F3D (7.16), 3F3E (7.70), 4MME (6.50) | Transporter |
| 52 | 2V7A (8.30), 3K5V (6.30), 3MSS (4.66), 3PYY (6.86), 4TWP (10.00) | Tyrosine-protein kinase ABL1 |

| ID | PDB codes in this cluster (with Binding Constant) | Protein Name |
|---|---|---|
| 53 | 3QGY (7.80), 4M0Y (6.46), 4M0Z (5.19), 4QD6 (8.64), 4RFM (10.05) | Tyrosine-protein kinase itk/tsk |
| 54 | 4E5W (7.66), 4IVB (8.72), 4IVC (10.00), 4IVD (9.52), 4K77 (6.63) | Tyrosine-protein kinase JAK1 |
| 55 | 4E6Q (8.36), 4F09 (6.70), 4GFM (7.22), 4HGE (7.92), 4JIA (9.22) | Tyrosine-protein kinase JAK2 |
| 56 | 1BZC (4.92), 2HB1 (3.80), 2QBP (8.40), 2QBQ (7.44), 2QBR (6.33) | Tyrosine-protein phosphatase non-receptor type 1 |
| 57 | 1C5Z (4.01), 1O5B (5.77), 1OWH (7.40), 1SQA (9.21), 3KGP (2.57) | Urokinase-type plasminogen activator |

**Supplementary Table 1:** CASF2016 Clusters
